# Exploring the immune environment of glioblastoma through single-cell RNA sequencing in humanized mouse models

**DOI:** 10.1101/2025.05.17.654526

**Authors:** Jun Takei, Ken Furudate, Yoshiko Nagaoka-Kamata, Opeyemi Iwaloye, Chloe E. Jepson, Madison T Blucas, Kiyotaka Saito, Robert S. Welner, Erwin G. Van Meir, Masakazu Kamata, Satoru Osuka

## Abstract

**Background:** Glioblastoma (GBM) is the deadliest primary brain tumor in adults, where current therapies fail to meaningfully extend survival. Available animal GBM tumor models, especially therapy-resistant and recurrent ones with unique immunological aspects, are restricted, impeding innovative treatment research. To confront this critical obstacle we established a unique GBM mouse model that utilizes patient-derived xenografts (PDXs) within humanized mice.

**Methods:** We selected two immune-deficient mouse models to facilitate the reconstitution of myeloid lineage cells. After undergoing myeloablation, mice received CD34+ hematopoietic stem progenitor cells derived from human umbilical cord blood for humanization. Upon confirming the reconstitution of human blood cells, mice were xenografted with PDXs resistant to radiation. Tumor profiles and immune cell infiltration were analyzed via flow cytometry, immunohistochemistry, and single-cell RNA sequencing (scRNA-seq). The findings were evaluated against scRNA-seq data from recurrent human GBM.

**Results:** A diverse range of human immune cells, including T, NK, and myeloid lineage cells, infiltrated PDX tumors in humanized mice. Notably, gene expression profiles in these immune cells resembled that of recurrent human GBM. Unlike conventional xenograft models, this model highlighted enhanced tumor diversity, particularly a high fraction of neural progenitor-like cells.

**Conclusions:** Our humanized GBM mouse model displayed an immune cell signature similar to recurrent GBM. This model is a valuable resource for analyzing the tumor immune landscape and assessing new therapies, particularly immunotherapies. By enabling effective evaluation of novel treatments, our model has the potential to significantly advance GBM research.

**Key Points:**

- A wide array of human immune cells infiltrates GBM-PDX tumors in humanized mice.
- The tumor heterogeneity of GBM-PDX is more evident in humanized mouse xenografts.

**Importance of the study:** Current GBM treatment development is primarily constrained by the absence of appropriate animal models that can enhance our understanding of human GBM biology and its immune dynamics. We created a specialized GBM mouse model using humanized mice with patient-derived xenografts resistant to radiation. Detailed immune cell analysis showed various human blood cells in the GBM-PDX derived from humanized mice. Single-cell RNA sequencing uncovered crucial T-cell subgroups, notably regulatory and exhausted T cells, which matched those observed in recurrent GBM patient tumors. Moreover, the model outperformed standard xenograft approaches in tumor heterogeneity, resembling the intricate diversity observed in recurrent GBM samples. Our humanized GBM mouse model provides a powerful platform to study the interactions between human tumors and immune cells. This tool will facilitate the discovery of immunosuppressive mechanisms and accelerate the assessment of immunotherapies applicable to various brain tumors.

## Introduction

Glioblastoma (GBM; 2021 WHO CNS grade 4) is the most frequent and lethal type of malignant brain tumor in adults, with an average survival of 15 months.^1,2^ Patients initially respond to standard therapies, including surgical excision, chemotherapy, and radiotherapy, but tumors recur and acquire resistance at a significant rate, leading to patient demise. In recent years, extensive genomic characterization of GBM has been performed, uncovering key oncogenic signaling pathways that drive the disease.^3,4^ However, strategies targeting these pathways have not significantly improved patient survival. Finding innovative approaches to lower mortality rates among GBM patients presents a significant challenge, which can only be overcome by enhancing our comprehension of the mechanisms that enable tumors to adapt to treatment and develop resistance.

GBM tumor tissue consists of a combination of neoplastic and stromal cells. The majority of non-neoplastic cells in tumors are immune cells, accounting for nearly 30% of the total volume,^5^ primarily consisting of microglia that reside in the brain and macrophages that originate in the bone marrow.^6,7^ Single-cell RNA-sequencing (scRNA seq) analyses of GBM tumors have shown that the immune populations present in the tumor microenvironment (TME) become more diverse upon tumor recurrence following radio- and chemotherapy.^7^ A higher percentage of T, natural killer (NK), and B cells are found in recurrent tumors.^7,8^ However, these immune cells fail to control tumor growth due to various tumor-related immune suppressive mechanisms, including the recruitment of regulatory T (Treg) cells, production of immunosuppressive cytokines, and up-regulation of checkpoint ligands.^9,10^ Developing effective treatments for GBM and assessing their *in vivo* effectiveness requires understanding the interaction between GBM cells and the immune system. However, the prevalent GBM tumor models consist of xenografting human cancer cells into immunocompromised mice or syngeneic mouse tumor cells in immunocompetent mice, neither of which are adequate for studying the human immune microenvironment in GBM.^11,12^

A humanized mouse model reflecting human immune cell dynamics has been established and is widely used to investigate tumor biology, immunology, and therapeutic interventions.^13,14^ This model is achieved by replacing the murine blood system with a human one after myeloablation and implantation of human hematopoietic stem progenitor cells (HSPCs) from various sources, such as umbilical cord blood (UCB), fetal liver, and peripheral blood (PB).^15,16^ Such models feature a blood system derived from humans, enabling investigations into human immune responses within a murine organism. They have become invaluable tools for the *in vivo* study of human hematopoiesis, immunology, and related diseases, including cancer. We have previously developed a human B-cell lymphoma humanized mouse model. It showed its use in assessing a novel anti-cancer treatment, including promoting antitumor memory responses.^13,14^ Diverse humanized mouse systems have also been utilized to examine the pathophysiology of GBM and to assess the effectiveness of treatment modalities using xenografted human GBM cell lines.^17–20^ These studies indicate that human GBM cells can generate tumors in humanized mice, which exhibit pathological attributes like those observed in patients with GBM. Flow cytometric assessments also indicated that various immune cell types, such as macrophages, CD4+ T, CD8+ T, and NK cells, had infiltrated within these tumors.^17^ However, the prior models utilized standard NSG or a related strain known as DRAG for humanization, which resulted in the suboptimal development of human myeloid lineage cells and Treg cells.^18^ The NOD.Cg-*Prkdc^scid^ Il2rg^tm1Wjl^*/SzJ (NSG) mouse strain has widely been used for mouse humanization. However, this strain is ineffective for human myeloid cell reconstitution because it lacks crucial human cytokines needed for myeloid lineage development.^13,14,21–24^ Myeloid cells are the most prominent immune cells in the patient’s GBM microenvironment, comprising 30-50% of the tumor mass.^25^ Several studies indicate that they render the tumor immunologically “cold”. Proficient myeloid lineage reconstitution is critical for adaptive immune responses^26,27^ as well as Treg cell differentiation via enhanced reconstitution of dendritic cells (DCs).^28,29^ Several new strains of mice were created to overcome the limitations of first-generation humanized mice.^30,31^ NSG-SGM3 (32) and NOG-EXL (33) second-generation models with an almost complete spectrum of human immune cells, except for microglial cells, are exciting. They still retain endogenous mouse CD45+ cells, but these are immature, and the mice lack a functional mouse immune system except for granulocytes.^32,33^ Both strains are derived from NOD-SCID (NOD.Cg-*Prkdc^scid^*) mice but differ in the extent of *IL2 common* γ *chain* (*IL2rg*) gene inactivation and the addition of human cytokine transgenes. NSG-SGM3 strain has a complete null mutation in *IL2rg*, while the NOG-EXL strain has only a partial deletion. The NSG-SGM3 expresses human IL3 (hIL3), GM-CSF (hGM-CSF), and SCF (hSCF), while the NOG-EXL expresses only hIL3 and hGM-CSF. Moreover, the NOG-EXL strain has nearly 10 times lower cytokine production compared to the NSG-SGM3, resulting in suppression of aberrant macrophage activation and exhaustion of HSPCs, enabling prolonged animal monitoring in research contexts.^31,34^

A thorough exploration of the immune cell presence in GBM tumors in these advanced humanized mouse models has not been performed. Furthermore, models of recurrent GBM, which exhibit a more profound immunosuppressive microenvironment, remain largely unexplored. To address this lack of knowledge, we examined the immune landscape of radio-resistant human GBM PDX xenografted in NSG-SGM3 and NOG-EXL humanized mice. We utilized immunohistochemistry, flow cytometry, and single-cell transcriptomic analysis to characterize and quantify the infiltrated immune populations. Our findings reveal that human GBM cells foster an immunosuppressive microenvironment, influence tumor cell behavior, and offer a new framework for testing therapeutic approaches.

## Materials and Methods

### Mouse humanization

Humanization of NSG-SGM3 and NOG-EXL mice was performed as reported previously.^21^ Before humanization with human CD34+ cells, 1-3 day old mice pups were irradiated with X-rays at 90cGy a day in advance. Purified CD34+ cells (0.1×10^6^ cells/mouse) were suspended in 20 µL of AutoMACS buffer (Miltenyi Biotec) and injected into the mice via facial vein as previously described.^35^ The levels of human blood cell reconstitution were monitored at 12-14 weeks post humanization.

### Single-cell RNA Sequencing (scRNA-seq)

The samples of PB, tumor-infiltrating lymphocytes (TILs), and tumor cells were obtained from NOG-EXL mice bearing EGFP+ JX-14P-RT cells. Following the manufacturer’s protocol, single-cell RNA libraries were prepared using the Chromium Single Cell 3’ Reagent Kits. Gene expression matrices were generated by the 10X Genomics CellRanger v6,1. The detailed methodology is described in the Supplementary Methods.

**Isolation of CD34+ cells from umbilical cord blood, Cell culture, Mice, Flow cytometry, Immunofluorescence, Public single-cell data analysis, and Statistical analysis.** These methods are described in Supplementary Methods.

## Results

### Humanized mice efficiently support GBM-PDX tumor growth

To inventory the array of human immune cells infiltrating GBM tumor tissues in a xenograft humanized mouse model, we used two independent immunodeficient mouse strains that support enhanced human myeloid lineage cell reconstitution compared to prior strains. The NSG-SGM3 strain has enhanced myeloid differentiation, while the NOG-EXL strain allows for extended monitoring of humanized mice post-tumor grafting.^34^ Humanized mice were prepared as previously reported (**Fig.1A**)^21^ (see detailed procedure in the Supplementaly Methods). The average frequency of humanization in mouse PB was 64.1% (48.0-94.5%) in NSG-SGM3 and 78.2% (62.9-97.0%) in NOG-EXL (p=0.0119, t-test) (**Fig.1B**), irrespective of donors used for humanization.

**Figure 1.**
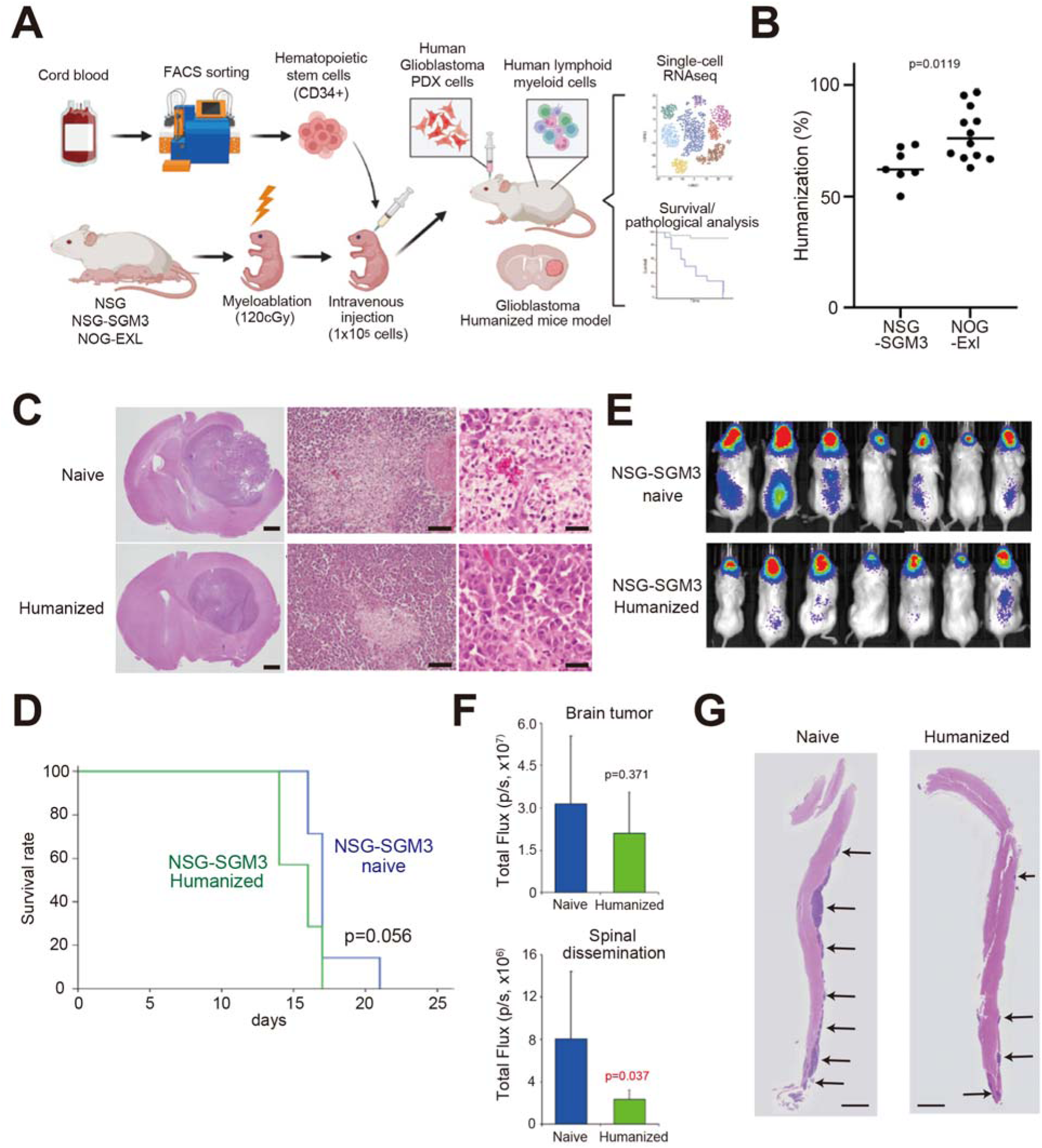
Human PDX GBM cells formed aggressive tumors in humanized mice. (A) Schematic showing the experimental design to establish humanized mice. Human HSCs were isolated from human cord blood. NSG or NSG-SGM3 newborn mice were irradiated with 120 cGy for successful engraftment. 1×10^5^ HSCs were injected intravenously. HSCs were engrafted in the mice, and the human immune system was reconstructed in the mice. HSCs differentiate into lymphoid and myeloid lineage cells. (B) The ratio of humanization in peripheral blood in humanized mice. (C) Histopathologic analysis of brain tumors in Naïve and Humanized mice. Scale bar: 1 mm (left), 100 μm (middle) or 50 μ m (right). (D) Survival curves for mice implanted with 4 x 10^5^ tumor cells (JX39P-RT) in NSG-SGM3 naïve and humanized mice. (7 mice/group; log-rank test). (E) Bioluminescence images of mice 2 weeks after tumor implantation. (F) Quantification data of (E) (Mann–Whitney U-test). (G) Histopathologic analysis of whole spinal cord in Naïve and Humanized mice. Disseminated tumor growth is identified in the area indicated by the arrows. Scale bar: 1mm.

We first evaluated the capability of humanized mice to support human GBM PDX growth in the presence of human myeloid lineage cell development using NSG-SGM3. This phase is vital for grasping human GBM biology in a humanized mouse model, particularly the mesenchymal subtype, where myeloid cells significantly contribute to tumor development.^36^ Flow cytometric analysis of PB confirmed that the NSG-SGM3 strain had stable engraftment of human myeloid lineage cells, including monocytes, macrophages, and granulocytes, in addition to NK cells, all of which failed to develop in non-humanized mice used as controls (**S-Fig.1**). As a source of human GBM, we selected the highly malignant human GBM PDX line, JX39P-RT, which recapitulates the characteristics of recurrent GBM. The JX39P-RT PDX line was developed from JX39P by subcutaneous implantation in nu/nu mice, followed by fractionated irradiation (2Gy x 6) and regrowth, repeated for 6 cycles.^37,38^ These cells harbor genetic alterations in EGFR vIII and CDK2NA, commonly observed in GBM.^39,40^ Following orthotopic xenografting into the right forebrain, the JX39P-RT cells formed aggressive tumors in both naïve and humanized NSG-SGM3 mice (**Fig.1C**). The tumors pathological examination revealed the pathognomonic features of human GBM,^41^ represented by pseudopalisading, hemorrhage, microvascular proliferation, necrosis, active proliferation, and cellular heterogeneity. Tumor histopathology was nearly identical between naïve and humanized mice. There was no statistically significant difference in survival between the xenografted naïve and humanized NSG-SGM3 mice (**Fig.1D**). JX39P-RT recapitulates the aggressive phenotype of recurrent GBM with a unique pattern of selective spinal cord metastasis following intracranial implantation. In contrast, intracranial tumor burden remained comparable, spinal cord dissemination was reduced four-fold in humanized mice (Figs.1E-F). H&E staining also confirmed that spinal dissemination was suppressed in the humanized mice (**Fig.1G**). The results indicate that the JX39P-RT PDX line can form aggressive intracranial GBM tumors in the severely immunocompromised NSG-SGM3 mouse strain, regardless of human immune cell reconstitution. However, the presence of human immune cells modulates tumor phenotype.

### A diverse range of human immune cells infiltrates GBM-PDX tumors in the brains of the xenografted humanized mice

The constrained distribution of GBM-PDX tumor cells in the spinal cord of humanized mice suggests that the involvement of human immune cells plays a role in the dynamics of GBM tumors. We next examined the interaction between human immune cells and GBM-PDX tumors by analyzing human CD45+ blood cells in the brains of humanized mice using flow cytometry. The human CD45+ blood cells in tumor tissue were analyzed against those in PB. Tumor samples from non-humanized naive mice were included as a negative control. Infiltration of various human immune cells was evidenced, which include CD4+ T, CD8+ T, CD19+ B, CD56+ NK cells, CD14+ monocytes/macrophages, and CD66b+ granulocytes (**Figs.2A-B,** n=7). While there was precise mouse-to-mouse variation, CD8+ T cells were the most abundant cell population, nearing 50% of total tumor-infiltrating human immune cells in 5/7 mice, and CD4+ T cells comprised a third of the cells in 4/7 tumors. The second significant population was monocytes/macrophages, which were present in all seven tumors at frequencies of ∼5-15%. Noteworthy, a substantial number of granulocytes and NK cells were found in the tumors. At the same time, B cells were also detected in lesser amounts (**Fig.2B**). A comparison of tumor-infiltrating immune cell populations versus PB evidenced great numbers in CD4+ T cells and B cells, as well as minor population of monocytes/macrophages and NK cells were seen in the blood. In contrast, high populations of CD8+ T cells, monocytes/macrophages, and CD56+ NK cells were identified in the tumors (**Fig.2C**), as observed in those with recurrent GBM clinical samples.^7,8^ Immunofluorescence (IF) staining of the tumor samples also confirmed the robust infiltration of various human immune cells into the tumors (**S-Fig.2**). These findings revealed that human immune cells within humanized mice could infiltrate GBM-PDX tumors.

**Figure 2.**
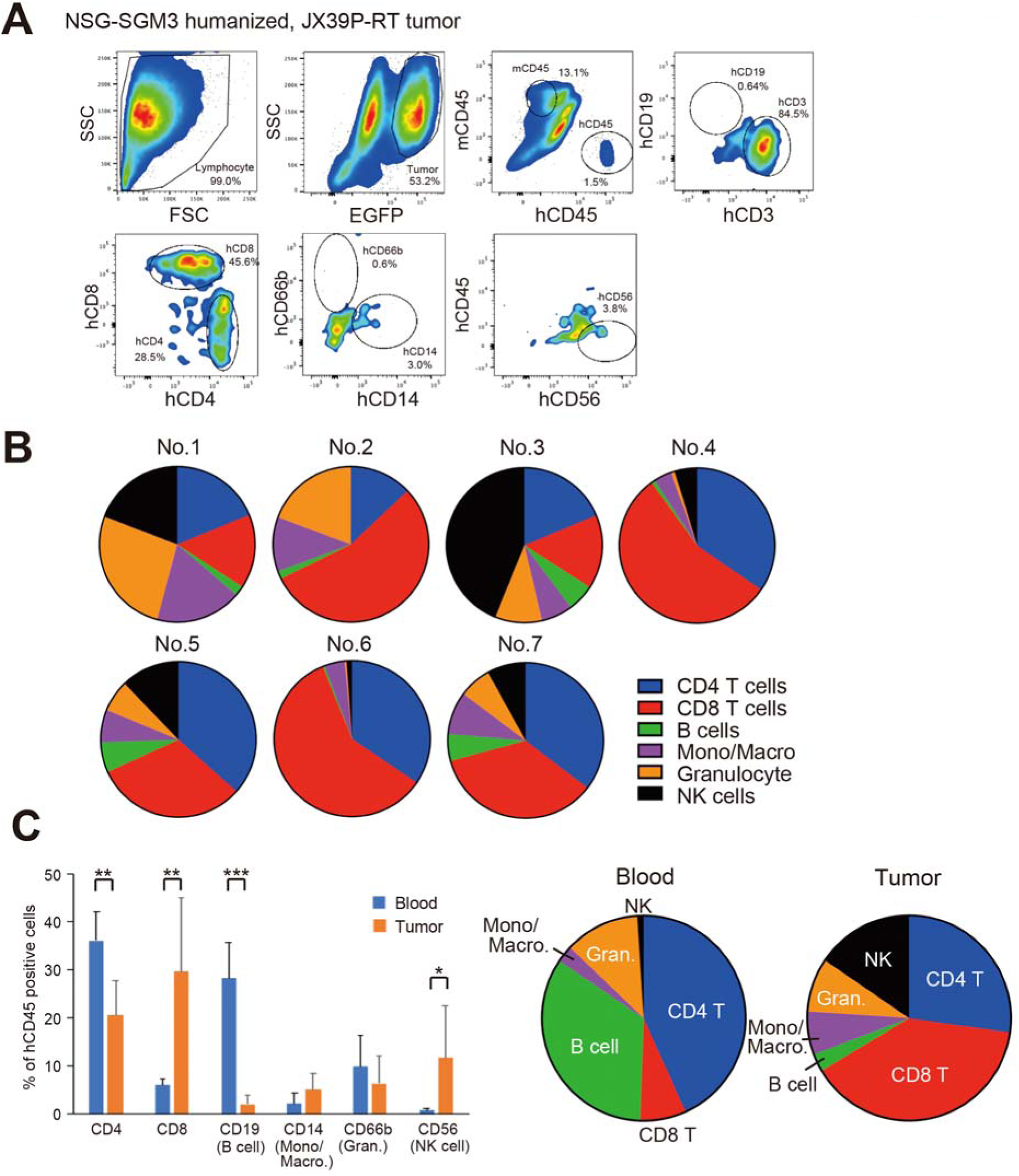
Human immune cells infiltrated into the brain tumors in humanized mice. (A) Representative data of Flow cytometry analysis for infiltrating immune cells in the brain tumors of humanized mice. %mouse CD45 (mCD45) and human CD45 (hCD45) in total cells. %hCD19, hCD3, hCD8, hCD4, hCD66b, hCD14 and hCD56 in hCD45 populations. (B) Percentage of immune cell types infiltrated within brain tumors. The following figure shows the findings in each of the seven Humanized mice. (C) (left) The comparison of the average ratio of several immune cells in peripheral blood and brain tumors. The ratio of human CD4, CD8, CD19, CD14, CD66b, and CD56 in hCD45 in blood and brain tumor. Two-tailed Student’s t-test. *P < 0.05, **P < 0.01, ***P < 0.001. (right) pie chart showing immune cell phenotypes in peripheral blood and GBM-PDX tumor of the tumor-bearing humanized mice.

### Single-cell RNA sequencing reveals diverse immune populations in the tumor microenvironment

The above findings highlighted robust TILs in the GBM-PDX tumors of NSG-SGM3 humanized mice. To better understand the immune landscape associated with these TILs, we conducted scRNA-seq on human CD45+ cell populations from peripheral blood leukocytes (PBLs) and GBM-PDX tumors grown in humanized mice. For these studies, we used another radiation-resistant recurrent GBM line, JX14P-RT,^37^ established from the JX14P PDX by repeated in vivo radiation exposure (2Gy x 6 x 6 cycles).^37^ These cells harbor genetic alterations in PTEN and CDK2NA, commonly observed in glioblastoma.^39,40^ The JX14P-RT cells exhibit slower tumor growth kinetics than those of JX39P-RT cells, allowing for more significant infiltration of human immune cells before animal demise. The median survival time for humanized mice bearing JX39P-RT PDX is 17 days, and 49 days for those with JX14P-RT PDX. We also utilized the NOG-EXL strain for humanization, which resulted in improved survival rates compared to the NSG-SGM3 strain, as previously reported.^34^ GBM-PDX tumor progression of JX14P-RT was also tracked via luciferase bioimaging. Human CD45+ TILs were isolated from three GBM-PDX tumor tissues 49 days post xenografting and subjected to scRNA-seq (**Figs.3-5**). 16,847 human CD45+ cells, consisting of TILs and PBLs, passed quality control. These cells were integrated to capture broad immune cell subsets, visualized using uniform manifold approximation and projection (UMAP) dimensionality reduction, resulting in 21 distinct clusters (**Figs.3A-B**). TILs displayed higher levels of several immune cell types compared to PBLs. Specifically, TILs had increased proportions of Tregs (cluster 0), CD8+ T cells (cluster 2), effector CD8+ cells (cluster 3), activated CD4+ cells (cluster 6), GZMK+ CD8+ cells (cluster 9), and memory CD4+ cells (cluster 10). Although not statistically significant, Tregs were more abundant in TILs than in PBLs (median 14.15% [interquartile range, 10.96–19.91%] vs. 8.34% [IQR, 6.75–8.88%]; p = 0.202) (**Figs.3B, 3C, and S-Table 1**) as known in patient GBM tumors.^42,43^ At the same time, there was a reduction in various populations of human PBLs, including naïve CD4+ and CD8+ T cells (clusters 4 and 7, p=0.00902 and p=0.0328), B cells (cluster 11, p=0.0163), and plasmacytoid DCs (clusters 17, p=0.0398) (**Figs.3B, 3C, and S-Table 1**). The overarching trends remained uniform despite observing differences in the donor mice’s blood cell population ratios (**Figs.3D-E and S-Figs.3A, 3B**). We also analyzed the TILs of GBM-PDX tumors compared to those obtained from the Extended GBmap, a public GBM scRNA-seq dataset (GSE211376).^44^ Immune cell proportions were extracted from 53 primary GBM cases and 26 recurrent GBM cases (**S-Table 2**). The results revealed that the proportions of CD4 and CD8 cells in CD45+ cells of humanized mice were higher than those found in recurrent GBM (p=0.0017), while the proportions of tumor-associated macrophages (TAMs) were found to be significantly lower (p=0.0498) (**Fig.3F**). The proportions of B cells, DCs and NK cells were similar in our model compared to human recurrent GBM (p=0.426, p=0.313, and p=0.359) (**Fig.3F**). Notably, microglia were absent in our humanized mouse models due to the use of HSPCs derived from UCB and a lack of human IL34 (**S-Table 2**).^45^ The above results reveal that both NSG-SGM3 and NOG-EXL humanized mouse models can replicate the TIL features associated with recurrent human GBM tumors.

**Figure 3:**
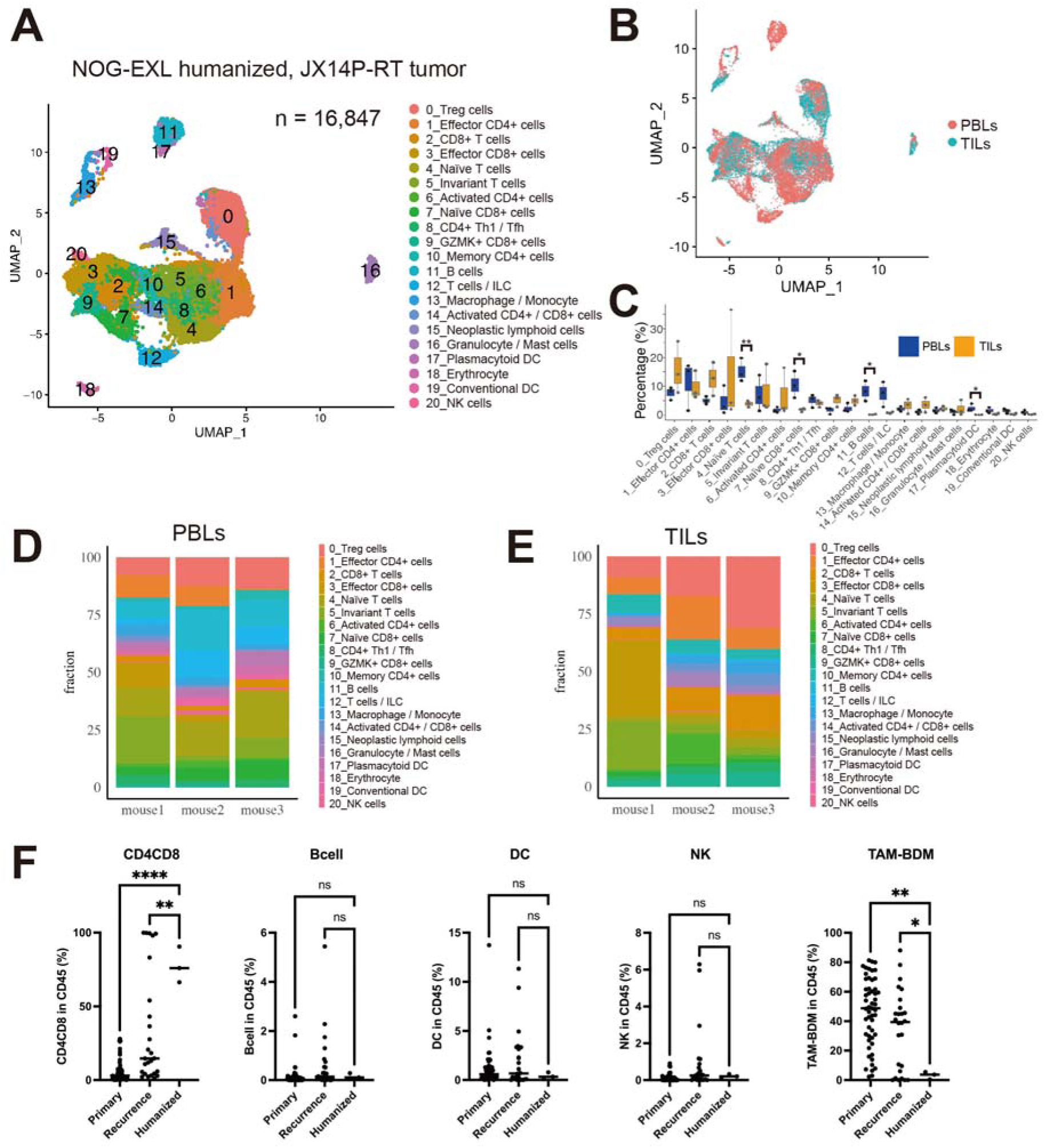
Single-cell RNA-seq analysis of JX14P-RT tumor in NOG-EXL humanized mouse model. (A) UMAP plot of cell types clustered by single-cell transcriptional analysis of CD45+ cells (n = 16,847 cells) from peripheral blood (PBLs) and tumor-infiltrating lymphocytes (TILs) isolated from NOG-EXL humanized mice bearing transplanted JX14P-RT human glioblastoma cells. (B) UMAP plot showing the origin of CD45+ cells as PBLs or TILs. (C) Comparison of cell type percentages between PBLs and TILs. Two-tailed Student’s t-test. *P < 0.05, **P < 0.01 (D) Distribution of 21 cell types in PBLs between three mice. (E) Distribution of 21 cell types in TILs between three mice. (F) Comparative analysis of tumor-infiltrating immune cell proportions within CD45+ populations across primary human glioblastomas (n = 53), recurrent human glioblastomas (n = 53), and tumors from humanized mice (n = 3). Human glioblastoma data were obtained from published datasets. Tumor associated macrophage – blood-derived monocyte/macrophage (TAM-BDM).

### T cells infiltrating GBM-PDX tumors are adopting immunosuppressive characteristics

To determine if the GBM-PDX humanized mouse model accurately reflects the immunosuppressive characteristic of recurrent patient GBM, we selected 8,650 CD3δ (CD3D)+ T cells, and the UMAP dimensionality reduction algorithm classified them into 13 clusters (**Fig.4A**) based on specific gene-expression pattern as reported previously.^46^ The identified clusters correspond to each CD4+ and CD8+ T cell subset, including naive, central memory (CM), effector memory (EM), effector (Effector), resident memory (RM), and Treg. The presence of two independent Treg populations (1 and 2) suggested that recurrent GBM may also harbor functionally diverse Treg populations in TILs, akin to previous findings in lung cancer.^47^ In addition, we observed clusters representing GZMK+ CD8+ T cells, which had a transcriptional program consistent with high expression of GZMK, found in the GBM TME.^46^ GZMK+ CD8+ T cells are a newly discovered immune cell population within the TME of GBM, characterized by relatively low expression of classical cytotoxic markers (PRF1, GLYN, and GZMB) and exhibiting an exhausted phenotype (**Figs.4B-C**). Although the specific function of GZMK+ CD8+ T cells remains ambiguous, their presence within GBM-PDX tumors is crucial for accurately interpreting the human GBM TME in a small animal model. Overall, the exhausted status of T cells was confirmed by the presence of four exhaustion-associated markers: CTLA4, PD1, LAG3, and TIGIT (**Fig.4D**). In Tregs (clusters 12 and 13), high expression of CTLA4 and TIGIT was observed. In contrast, CD8+ T cells (clusters 7, 8, 9, 10, and 11) showed high expression of PD-1 and LAG3, which is consistent with a previous report.^48^

**Figure 4:**
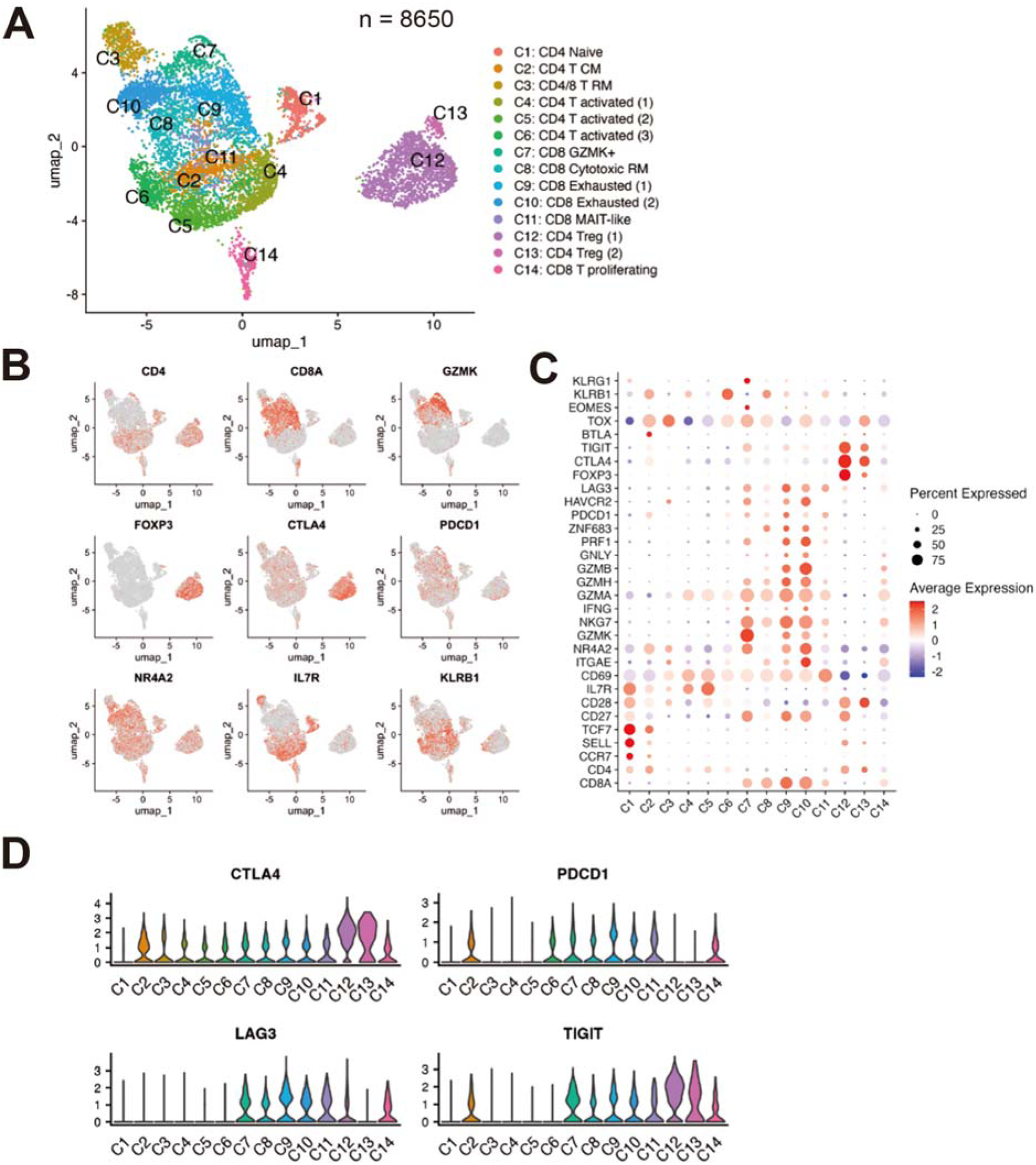
Detailed characterization of T cell subsets infiltrating brain tumors in humanized mice model. (A) UMAP plot of T cell populations (n = 8,650 cells) from JX14P-RT tumors (n = 3) in NOG-EXL humanized mouse model showing 14 distinct cell types identified. (B) Feature plot showing the expression of key T cell markers across the identified subsets. The intensity of color represents the level of gene expression for KLRB1, PDCD1 (PD-1), IL7R, CTLA4, CD8A, NR4A2, FOXP3, CD4, and GZMK. (C) Heatmap displaying the expression patterns of T cell-associated genes across the 13 identified subsets. (D) Dot plot showing the expression of four key immune checkpoint molecules (CTLA4, TIGIT, PD1, and LAG3) across the T cell subsets.

To determine whether the CD3δ+ T cell composition in GBM-PDX tumors mirrors that found in patient-derived GBM tumors, we analyzed scRNA-seq data from CD3δ+ T cells alongside the recurrent GBM dataset.^44^ A total of 44,560 cells were extracted and visualized using UMAP dimensionality reduction (**S-Fig.4A**). Cluster-defining genes and select functional markers are shown in **S-Figs.4B, 4C**. Within the TME of recurrent GBM, clusters of CD4+ T CM, CD4+ Treg, CD8+ GZMK+, CD8+ cytotoxic RM, and CD8+ exhausted T cells were observed, showing commonality with the TME of humanized mice. Findings from the comparative analysis indicate that the TME of GBM-PDX tumors derived from humanized mice closely reflects the characteristics of recurrent GBM tumors, particularly regarding T cell subset distribution and gene expression patterns.

### The humanized mouse model demonstrated extensive Treg cell infiltration

Treg cells are critical in establishing immunosuppressive TME in recurrent GBM.^49–51^ We thus conducted an in-depth study of Treg cells within GBM-PDX tumors from the humanized mice. 1,528 Treg cells were identified among 8,650 CD3δ+ T cells within the tumors. These Tregs cells were classified by the UMAP dimensionality reduction algorithm into five distinct clusters, including four “activated” clusters (C1-C4, **Fig.5A**) and one resting cluster (C5, **Fig.5A**) classified by high expression of IL2RA, CCR8, MAGEH1, and LAG3, which are associated with Treg activation (**Figs.5B-C**).^47,52^ The first subcluster (C1/Activated 1/OX40^hi^GITR^hi^) exhibited a gene expression profile indicative of a distinct activated state. This was characterized by elevated expression levels of tumor necrosis factor receptor superfamily (TNFRSF) 4 (also known as OX40) and TNFRSF18 (also known as GITR) (**Figs.5B-C and S-Fig.5**).^53^ The expression levels of additional genes related to Treg activation increased, with some genes being prevalent across multiple activated clusters; LAG3 (C2/Activated 2), ICOS, and TNFRSF9/4-1BB (C3/Activated 3, and IKZF2/Helios) (C4/Activated 4). To assess the activation status of Tregs, we analyzed additional activation-associated markers, such as CTLA4, LAG3, TNFRSF18, and MAGEH1 (**Fig.5D**).^52,53^ In all five clusters, high expressions of CTLA4 and TNFRSF18 were observed. Additionally, activated clusters C1-3 showed high expression of LAG3 and MAGEH1, indicating that these cells have tumor-effector Treg phenotypes.^52^ However, activated C4 showed low expression levels of LAG3 and MAGEH1 compared to other activated clusters, suggesting that this cluster may represent peripheral effector Tregs (**Fig.5D**).^52^ These results indicate that GBM-PDX tumors harbor heterogeneous Tregs with diverse phenotypes, each exhibiting distinct immunosuppressive functions.^47,54,55^

**Figure 5:**
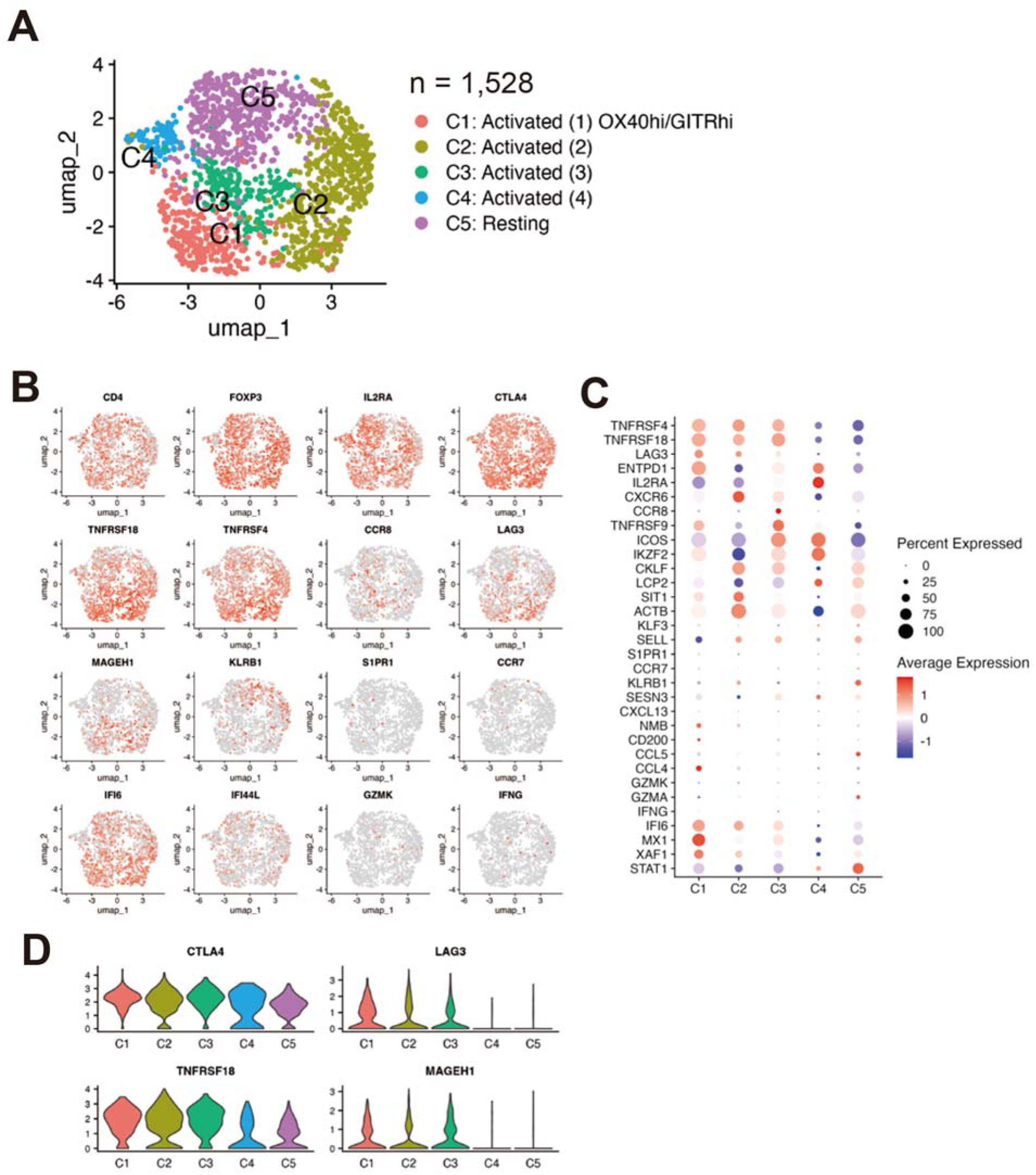
In-depth analysis of regulatory T cell (Treg) subpopulations isolating from brain tumors in humanized mice model. (A) UMAP plot of Treg cells (n = 1,528) from JX14P-RT tumors (n = 3) in NOG-EXL humanized mouse model showing five cell types. (B) Feature plots displaying the expression of individual genes across the Treg population. (C) Heatmap showing differential gene expression across the identified Treg subsets. (D) Violin plot showing the expression of four key immune checkpoint molecules (CTLA4, TIGIT, PD1, and LAG3) across the T cell subsets.

We further assessed the above outcomes using the human recurrent GBM dataset,^44^ following the same approach as performed in **Fig.4**. A total of 2,661 cells were extracted and visualized using UMAP dimensionality reduction (**S-Figs.5A-B**). We identified five unique clusters in this dataset, differing slightly from those in GBM-PDX tumors. In the ‘activated’ clusters, activation genes were expressed, and some showed exceptionally high expression across different activated clusters; OX40 and GITR (Activated 1_OX40hi GITRhi), LAG3 (Activated 2), and TNFRSF9 (Activated 3) (**S-Figs.5C-D**). In all five clusters, the high expression of CTLA4 and TNFRSF18 was observed; however, activated clusters (clusters 1-3) exhibited the high expression of MAGEH1 (**S-Fig.5E**). Importantly, the activation status of Tregs resembled that found in the GBM-PDX tumors from xenograft humanized mice. In all five clusters, the high expression of CTLA4 and TNFRSF18 was confirmed; however, activated clusters (clusters 1-3) also showed the high expression of MAGEH1 (**S-Fig.5E**). As mentioned above, these in-depth comparative analyses demonstrated that the humanized mouse model closely recapitulates the complex Treg profile observed in recurrent human GBM.

### The humanized mouse model demonstrated MDSC infiltration

Myeloid-derived suppressor cells (MDSC) play a crucial role in sustaining immunosuppression in the TME by supporting the recruitment and inhibitory functions of Tregs.^56^ We thus investigated MDSC populations in humanized mice. A total of 321 macrophages and monocytes were identified and classified by the UMAP dimensionality reduction algorithm into four distinct clusters (C1-C4, **S-Fig.6A**), based on the expression levels of CD14, FCGR3A, S100A8, S100A9, and MRC1 (CD206), which are associated with the immunosuppressive functions of myeloid cells (**S-Figs.6B-C**).^7,57^ The first subcluster (C1/Monocytic MDSC) exhibited a gene expression profile indicative of an immunosuppressive state. This was characterized by high expression levels of CD14, S100A8, and MRC1 (**S-Figs.6B-C**).^53^ In all four clusters, high expression of S100A9 was observed, suggesting that these macrophages (C2-C4) were differentiated from M-MDSCs, but not from monocytes (**S-Fig.6D**).^57^

Using the same approach as performed in **Fig.4**, a total of 19,414 cells were extracted and visualized using UMAP dimensionality reduction from the human recurrent GBM dataset^44^ (**S-Figs.7A**). We identified 16 clusters, which were more abundant than clusters in GBM-PDX tumors. Interestingly, we identified two MDSC clusters (C5 and C6) and two M2-like macrophage clusters (C15 and C16), which revealed high expression levels of S100A9 and MRC1 (**S-Fig.7B-D**). These in-depth comparative analyses demonstrated that the humanized mouse model recapitulates the MDSC and macrophages, which exhibit the immunosuppressive phenotypes observed in recurrent human GBM.

### The human immune microenvironment plays a crucial role in preserving the diversity of tumor characteristics in GBM-PDX tumors

The above findings indicate that the GBM-PDX tumors obtained from xenografted humanized mice reflect a detailed TIL profile similar to that found in recurrent human GBM from patients. One of the primary traits of recurring human GBM is the marked diversity observed within the tumor, which contributes significantly to treatment resistance.^58–60^ However, achieving an accurate representation of tumor diversity in a standard GBM-PDX mouse model remains a significant challenge.^12^ Finally, we conducted a comparative study for GBM-PDX tumor cells in the absence (i.e., naïve mice) and the presence of the human blood system (i.e., humanized mice). Using EGFP fluorescence as a marker, JX14P-RT cells were extracted from three humanized and two naïve mice brains. 6,180 cells from xenograft humanized mice and 5,516 cells from xenograft naïve mice were identified, respectively, and classified by the UMAP dimensionality reduction algorithm into 17 clusters (**Fig.6A**). A distinct separation on the UMAP plot was confirmed between tumor cells from humanized mice and naive mice (**Fig.6B**), and this separation pattern was observed in multiple independent samples (**Fig.6C**). To quantify tumor cell heterogeneity, we calculated the Euclidean distance of individual cells from their cluster centers in UMAP space, providing a quantitative measure of cellular dispersion and heterogeneity. Statistical analysis revealed significantly higher distance values in cells from humanized mice compared to non-humanized mice (Wilcoxon test, p < 0.001) (**Fig.6D**). These results indicate heightened tumor variability in the humanized mouse model.

**Figure 6:**
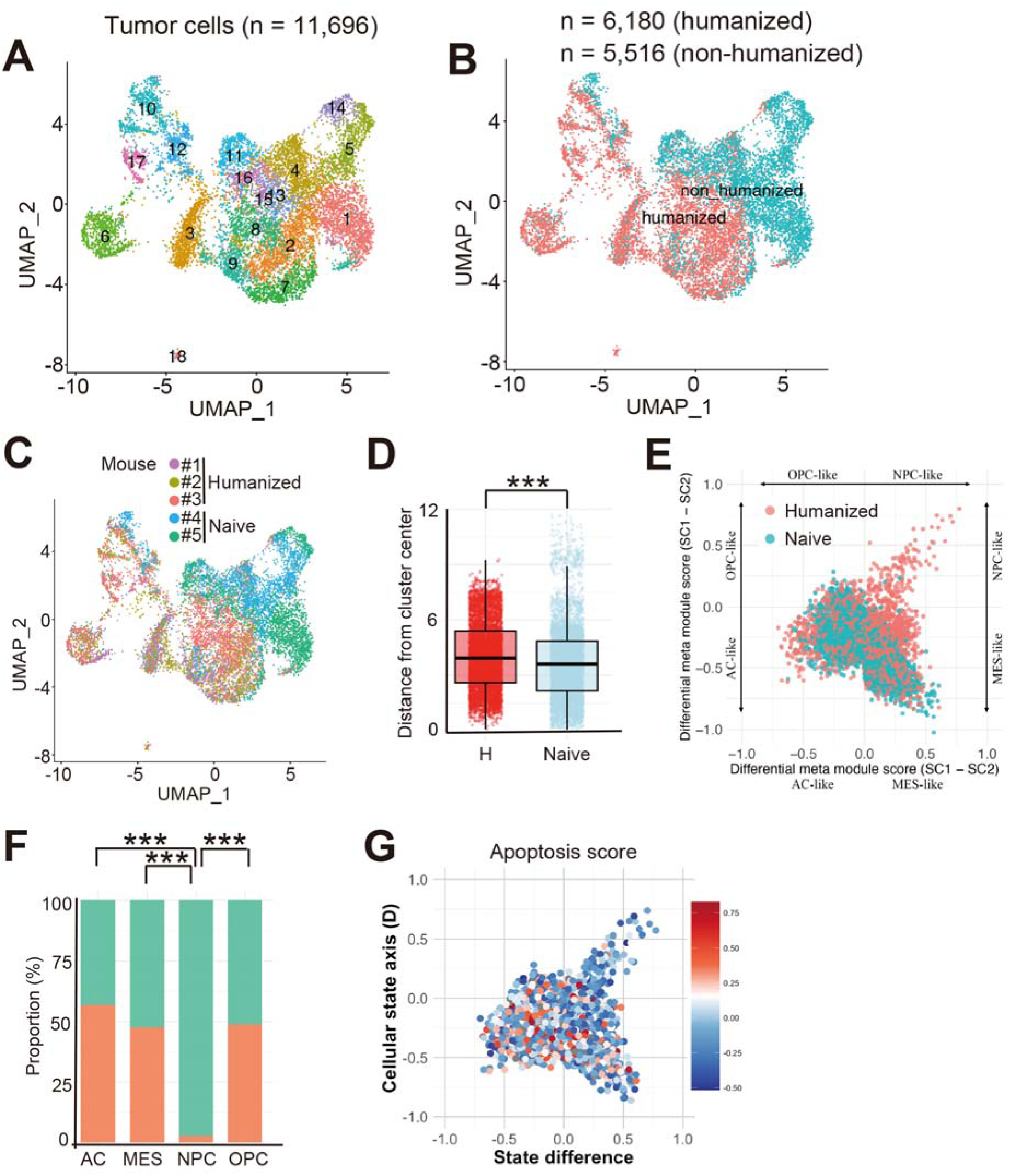
Comparison of brain tumor characteristics between humanized and naive mouse models. (A) UMAP plots of tumor cells from individual humanized mice (n=3) and naive mice (n=2), highlighting 17 distinct cell types. (B) UMAP plots illustrate differences between humanized and naive mouse tumors. (C) UMAP plots showing tumor cell distribution for each humanized and naive mouse. (D) Distribution of tumor cell distances from cluster centers in UMAP space comparing humanized (n = 6,180) and naive mice (n = 5,516). Wilcoxon test, ***P < 0.001. (E) Two-dimensional representation of cellular states where four regions indicate NPC, OPC, AC, and MES states. Each dot represents the exact position of individual tumor cells, reflecting their relative meta-module scores. Dot colors distinguish between tumor cells derived from humanized or naive mice. (F) Proportional distribution of tumor cell origins in astrocyte (AC), mesenchymal (MES), neural-progenitor (NPC), and oligodendrocyte-progenitor (OPC) states. Kruskal-Wallis rank sum test, ***P < 0.001. (G) Distribution of apoptosis scores across four distinct cellular states. The color intensity represents the apoptosis score of individual cells within each state.

Tumor diversity is heavily influenced by how tumor cells are distributed throughout the cell cycle.^61^ The annotation of cell cycle genes revealed an increase in S-phase cells (p = 0.0048) and a reduction in G2/M-phase cells (p = 0.001) in the GBM-PDX tumors from humanized mice (**S-Fig.8A**). These findings were consistently observed across different samples (**S-Fig.8B**).

### Human GBM PDX tumor cells exhibit increased stemness under the human immune microenvironment

In GBM, elevated stemness properties correlate with stronger resistance to radiotherapy (and chemotherapy) and a greater capacity to evade immune responses, notably in recurrent GBM.^62–64^ We thus examined the differentiation state of GBM-PDX tumors using the computational analysis methods described by Neftel et al.^65^ This strategy divides GBM cells into four separate differentiation categories: neural-progenitor (NPC), oligodendrocyte-progenitor (OPC), astrocyte (AC), and mesenchymal (MES). The classification is based on the expression profiles of markers that are characteristic of various stages seen in patient-derived GBM tumors. Strikingly, GBM-PDX tumors exhibited distinct alterations, most notably a significant increase in the proportion of NPC-like cell fraction, marked by enhanced expression of SOX4 and Tubb3 (**Figs.6E-F**).^65^ The other three populations showed no significant discrepancies. These findings were confirmed in three humanized mice models and two non-humanized mice models used as controls (**S-Fig.9A**). We confirmed that the markers for the four clusters are elevated in their respective clusters (**S-Figs.9B-C**). The increased levels of SOX4 expression were confirmed in a different GBM-PDX humanized model, specifically JX39P-RT (**S-Fig.9D**).

The presence of NPC-like cells in GBM-PDX tumors might reflect a significant evolutionary response to immune pressure for survival. To verify this hypothesis, we evaluated the extent of apoptosis among the four tumor cell populations using an apoptosis score calculated based on the expression levels of canonical apoptosis markers.^66,67^ As expected, the NPC-like cell population exhibited lower apoptosis scores compared to other populations (**Fig.6G and S-Fig.9E**). The data convincingly demonstrate that GBM-PDX tumors arising in humanized xenograft mice can support the presence of cells with NPC-like cell phenotype. In contrast, this phenotype was not observed in immunodeficient mouse models. In summary, incorporating a human immune environment is crucial in generating a GBM-PDX tumor phenotype that authentically represents the recurrent GBM tumors observed in clinical samples.

## Discussion

Using animal models in cancer research is essential for bridging the gap between preclinical studies of human cancer biology and evaluating new therapeutic strategies’ efficacy. In human GBM research, it is common to use immune-compromised mice grafted with human PDX GBM. Recent advancements in human cancer animal models using humanized mice have provided opportunities to study the dynamics between human cancers and immune cells in the complex TME.^13,14^ In this study, we developed a humanized mouse model that incorporates human GBM PDX tumors with radiation resistance, enabling a detailed examination of recurrent GBM in a nearly complete human immune microenvironment. Our model’s significant improvement lies in using advanced NSG strains, including NSG-SGM3 and NOG-EXL. Both models enhance myeloid lineage human cell reconstitution, including MDSCs,^68,69^ a process inadequately supported by standard humanized NSG mice.^21^ Utilizing these models resulted in three primary insights; (1) these humanized mouse models reflected the immune characteristics of recurrent GBM, evidenced by the infiltration of various human immune cells and transcriptional similarities with tumors from patients, (2) there was marked infiltration of Treg cells exhibiting activation features similar to the immunosuppressive behavior seen in recurrent human GBM, and (3) the presence of immune cells in humans enhanced tumor variability and stemness in GBM PDX cells, notably fostering therapy-resistant NPC-like cell development. These results indicated that GBM PDX mice with a human immune microenvironment effectively mimicked the conditions in human patients, revealing immune responses similar to those in recurrent human GBM tumors.

T-cell responses play a crucial role in tumor immunity, and their dysfunction is a hallmark of recurrent GBM.^70,71^ Prior studies have demonstrated that the GBM TME is characterized by infiltration of Tregs and exhausted T cells, which contribute to local immunosuppression and treatment resistance.^70,72,73^ The humanized mouse models we created, utilizing human GBM PDX with radiation resistance, effectively replicated the key T cell response traits observed in recurrent GBM patients. While studies involving humanized mice indicate that human T cells can infiltrate GBM tumor xenografts, there has been limited in-depth examination of these T cells. The primary reason for this limitation was the inadequate restoration of MDSCs. MDSCs are indispensable for cultivating a suppressive environment in GBM and are significantly associated with the induction of T-cell exhaustion.^74–76^ In addition, MDSCs contribute to the recruitment and subsequent activation of Tregs in the TME through both direct and indirect mechanisms.^69,75,77–79^ Through scRNA-seq analysis, we conducted the first comprehensive examination of detailed T-cell populations in GBM humanized mouse models. Our study revealed remarkable similarities between the T cell landscape in our model and that observed in human recurrent GBM. We identified distinct populations of activated Tregs expressing high levels of OX40, GITR, and other activation markers, closely matching the phenotypes found in patient tumors. The presence of GZMK+ CD8+ T cells, a recently identified population in human GBM showing characteristics of T cell exhaustion, further validates the physiological relevance of our model. Another significant finding was the identification of heterogeneous Treg populations within the TME, including multiple activated subsets with distinct molecular signatures.^51,80^ The presence of these diverse Treg populations and their expression of key immunosuppressive markers suggests that our model effectively reproduces the complex immunoregulatory networks present in human GBM. Furthermore, the observation of exhausted T cells, especially in CD8+ T cells, expressing multiple checkpoint molecules indicates that our model captures the T cell dysfunction characteristic of recurrent GBM.

A hallmark of recurrent GBM is the emergence of therapy-resistant tumor cell populations that arise through complex interactions with the TME. These adaptations include increased stemness, altered cell cycle dynamics, and enhanced immune evasion capabilities. The scRNA-seq study indicated that the human immune environment in GBM-PDX mice exhibited significant phenotypic variations, corresponding to key features of recurrent human GBM tumors. There was a notable rise in tumor diversity and the presence of tumor cells with enhanced stem cell-like features, which were unique to the humanized mice and not found in standard immunodeficient mice. This unexpected result highlighted the distinct advantages of our humanized mouse model in exploring interactions between GBM and the immune system. As a result, this model establishes a strong basis for delving into the detailed mechanisms that contribute to immune evasion and therapy resistance in GBM. Furthermore, this approach provides a critical resource for assessing novel therapies, especially those that exhibit strong interactions with the immune systems of hosts, to determine their efficacy.

Our humanized mouse model effectively mirrors numerous phenotypic characteristics of human GBM tumors when integrated with the human immune system; however, future research must tackle several identified limitations. A key finding was the lower percentage of tumor-associated macrophages than human GBM tumors. While the observations could be unique to the GBM cell lines we tested, it is vital to further research with various GBM models to comprehensively understand the factors that drive macrophage infiltration and their functional contributions in our experimental setup. Another major shortcoming of our present model is the exclusion of human microglial populations. The significance of human microglia in GBM tumor biology is clear,^81^ however, the reconstitution of human microglia in mice is challenging since they derive from primitive myeloid progenitors in the yolk sac that are not ideal for research.^82,83^ Recently, a team led by Gorantla successfully engineered a new humanized NOG mouse model, enabling the derivation of microglia through the transgenic expression of human interleukin-34 (hIL34).^45^ Although the mouse cannot regulate hIL34 levels due to the systemic expression from the transgene, employing this strain in a GBM-PDX humanized mouse model would enhance our understanding of microglial involvement in tumor biology and the assessment of antitumor therapies. From a different perspective, the systemic production of hIL34 triggers the abnormal activation of cells responsive to IL34, notably monocyte-macrophage lineage cells and Treg cells.^84,85^ Moreover, our current model uses HLA-mismatched combinations of tumor cells and immune cells, which could potentially trigger unintended immune responses that may not reflect the natural course of GBM progression. Future studies should consider developing models that use patient-matched HSPCs and tumor cells to address this limitation. Such matched models would more accurately represent the immune responses occurring in GBM patients and offer a more reliable platform for evaluating immunotherapeutic strategies.

## Required Statements

### Ethics

The collection of human umbilical cord blood (UCB) samples was approved under IRB protocol (IRB-300004736) at the University of Alabama at Birmingham Hospital. Animal research described in the study was approved by the University of Alabama at Birmingham (UAB) ’s Chancellor’s Animal Research Committee (Institutional Animal Care and Use Committee [IACUC]) and was conducted in accordance with guidelines for housing and care of laboratory animals of the National Institutes of Health and the Association for the Assessment and Accreditation of Laboratory Animal Care International.

### Authorship

JT, KF, OI, RSW, and SO performed scRNA-seq analysis. JT and SO performed an an immunohistochemistry and immunofluorescence analysis. YNK, MTB, SUA, and CEJ prepared humanized mice and maintenance of them. OI, KS, and SO performed stereotactic neurosurgery. SO performed an analysis of the tumor tissue. SO, EGVM and MK conceived the project, designed experiments, and wrote the manuscript.

### Data Availability

Prior to manuscript publication, we will deposit our Single-cell RNA sequencing data in the Gene Expression Omnibus (GEO) repository for public access.

## Acknowledgments

We thank all members of our laboratories for their helpful advice. We appreciate the help from the UAB Flow Cytometry and Single Cell and High-Resolution Imaging shared resources and the Center for AIDS Research grant AI027767. Illustrations were created with BioRender (https://biorender.com/). We gratefully acknowledge the provision of supercomputing resources by the Human Genome Center of the University of Tokyo’s Institute of Medical Science. We acknowledge the use of AI-assisted language tools, including ChatGPT and Claude, for English editing and proofreading of this manuscript.

## Abbreviations

GBM: Glioblastoma
TME: tumor microenvironment
scRNA-seq: single-cell RNA sequencing
PDX: patient-derived xenograft
HSPC: hematopoietic stem progenitor cells
NSG: NOD.Cg-*Prkdc^scid^ Il2rg^tm1Wjl^*/SzJ
NSG-SGM3: NOD.Cg-*Prkdc^scid^Il2rg^tm1Wjl^*Tg(CMV-IL3,CSF2,KITLG)1Eav/MloySzJ
HSPCs: hematopoietic stem progenitor cells

