## Supplementary materials for "Exploring the immune environment of glioblastoma through single-cell RNA sequencing in humanized mouse models"

**Supplementary documents**

**Supplementary methods**

**Isolation of CD34+ cells from umbilical cord blood**

All human umbilical cord blood (UCB) samples were collected from females following vaginal delivery at the University of Alabama at Birmingham Hospital, giving written informed consent under an approved IRB protocol (IRB-300004736). UCB was collected to a sterile cord blood collection unit (MSC127D, MacoPharma, GA) and processed within 24 hours as described before.^1^ Briefly, the mononucleated cell (MNC) fraction was separated by Ficoll-Paque density gradient centrifugation, followed by CD34+ cells in the MNC fraction were positively isolated using CD34 beads (130-046-703, Miltenyi Biotec, Germany) via autoMACS Pro Cell Separator (Miltenyi Biotec) according to the manufacturer’s protocol. Obtained CD34+ cells were cryopreserved in liquid nitrogen until use.

**Cell culture**

JX39P-RT and JX14P-RT are radio-resistant GBM patient-derived xenografts (PDX) cells generated by Dr. Christopher Willey (UAB) as previously described.^2,3^ Briefly, primary PDX was established by growing patient-derived GBM cells subcutaneously in nude mice. Subcutaneous tumors of JX39P or JX14P were exposed to fractionated irradiation (2 Gy x 6, 3 times per week for 2 weeks), and then recurrent tumors were passed to a new set of mice. The same protocol of irradiation and passage was performed 6 times to establish radio-resistant PDX (JX39P-RT and JX14P-RT). Human Glioma stem cells from primary and radio-resistant PDX tumors were established and cultured in a Neural stem cell medium.^2,4^

**Mice**

NSG-SGM3 [NOD.Cg-*Prkdc^scid^Il2rg^tm1Wjl^*Tg(CMVIL3, CSF2, KITLG)1Eav/MloySzJ)] mice were purchased from the Jackson Laboratory (#013062). NOG-EXL (NOD.Cg-*Prkdc^scid^ Il2rg^tm1Sug^*Tg(SV40/HTLV-IL3,CSF2)10-7Jic/JicTac) mice were sourced from Taconic Biosciences (#13395). Orthotopic implantation of cells was performed as described.^2,4,5^ Briefly, viable GBM patient-derived xenografts (PDX) cells were injected into the right forebrain with 400,000 cells for JX39P-RT and 200,000 cells for JX14P-RT into NSG-SGM3 and NOG-EXL mice, respectively. Tumor growth was monitored via luciferase bioimaging using an IVIS III bioluminescence imager (PerkinElmer) as previously described.^2^

**Mouse humanization**

Humanization of NSG-SGM3 and NOG-EXL mice was performed as reported previously.^1^ Before humanization with human CD34+ cells, 1-3 day old mice pups were irradiated with X-rays at 90cGy a day in advance. Purified CD34+ cells (0.1x10^6^ cells/mouse) were suspended in 20 µL of AutoMACS buffer (Miltenyi Biotec) and injected into the mice via facial vein as previously described.^6^ The levels of human blood cell reconstitution were monitored at 12-14 weeks post humanization. Littermates of the same sex were randomly assigned to experimental groups. Sample sizes in each experiment are included in legends.

**Flow cytometry**

Flow cytometry was performed on a BD FACSymphony (BD medical device, NJ) using a gating strategy as described previously^1^ and analyzed using FlowJo 10 (FlowJo, LLC, OR). Briefly, 100 µL of peripheral blood was collected from humanized mice via retro-orbital vein at 14 weeks of age following humanization as well as from xenograft mice at the endpoint. The blood samples were stained with antibodies against mouse CD45, human CD45, CD3, CD4, CD8, CD14, CD19, CD56, and CD66b 30 min at 4 °C and fixed in RBC Fix & Lysis (422401, BioLegend, CA), making the total volume 1000 µL. Tumor-infiltrating human immune cells were obtained according to a protocol reported previously.^7^ Briefly, mice were first perfused with PBS to remove all circulating blood cells, and then brain tumor tissues were dissected and minced into 0.5-1 mm^3^ pieces. The tissues were then incubated with Liberase TL (Sigma-Aldrich), which contains negligible levels of endotoxin together with DNase 1 (TURBO DNase, Thermo Fisher Scientific). It is highly important to maintain isolated immune cells without unexpected activation.^8^ Any undigested debris was removed through a 30 µm MACS SmartStrainer (Milteny Biotec) prior to staining. All antibodies were obtained from BioLegend: APC anti-human CD66b (396906, clone QA17A51), PE/Cyanine7 anti-human CD3 (344816, clone SK7), Brilliant Violet (BV) 421 anti-human CD56 (362552, clone 5.1H11), BV510 anti-human CD4 (357420, clone A161A1), BV605 anti-human CD8 (344742, clone SK1), BV711 anti-human CD19 (302246, clone HIB19), BV785 anti-human CD14 (301840, clone M5E2), BV650 anti-human CD45 for all human leukocytes (304044, clone HI30), and APC/Fire750 anti-mouse CD45 for all mouse leukocytes (147714, clone I3/2.3). EGFP positive fraction for GBM PDX tumor cells, as well as APC/Fire750 positive fraction for mouse leukocytes, were gated out for analysis of the BV650 positive human leukocyte population.

**Immunofluorescence**

Mouse brain tumors were fixed by 4% paraformaldehyde overnight, and then permeabilized with 1% Triton X-100 for 10 minutes. Tissues were blocked with 3% BSA in PBS for 1 hour and incubated overnight with an anti-CD4 antibody (Invitrogen, 14-0049-82), anti-CD8 antibody (Invitrogen, MA8-14348), anti-CD56 antibody (Invitrogen MA-1-06801), Anti-CD68 antibody (Invitrogen, MA5-13324), Anti-Collagen type 4 antibody (Millipore, AB756P) or Anti-Glut1 antibody (Millipore, 07-1401). After washing with PBS, tissues were cultured overnight with Alexa Fluor 594 goat anti-Rabbit IgG (H+L) (Fisher, A11037), Alexa Fluor 594 donkey anti-Mouse IgG (H+L) (Fisher, A21203), Alexa Fluor 647 goat anti-Rabbit IgG (H+L) (Fisher, A21245) or Alexa Fluor 647 goat anti-Mouse IgG (H+L) (Fisher, A21236). Images were taken by using a confocal microscope (Nikon, A1R). Nuclei were counterstained with Hoechst 33342 (Invitrogen, H3570).

**Single-cell RNA Sequencing (scRNA-seq)**

The samples of peripheral blood (PB), tumor-infiltrating lymphocytes (TIL), and tumor cells were obtained from NOG-EXL mice bearing EGFP+ JX-14P-RT cells. Single-cell suspensions were prepared by using the Brain Tumor Dissociation Kit and gentleMACS™ Dissociators (Milteny Biotec). CD45+ cells and EGFP + cells from tumor tissues were extracted using a cell sorter (ARIA I). CD45+ cells in PB were purified using EasySep Mouse/Human Chimera Isolation Kit (STEMCELL). Cell viability was assessed using trypan blue exclusion; only samples with >90% viability were used. Surface staining was performed using TotalSeq™-B anti-human hashtag antibodies described in the BioLegend protocol (TotalSeq™-b or -c with 10x feature barcoding technology. Available at: <https://www.biolegend.com/fr-ch/protocols/totalseq-b-or-c-with-10x-feature-barcoding-technology>). Following the manufacturer’s protocol, single-cell RNA libraries were prepared using the Chromium Single Cell 3’ Reagent Kits. Briefly, 20,000 cells were loaded onto the 10x Genomics Chromium to capture approximately 20,000 cells. Reverse transcription and cDNA amplification were performed according to the kit protocol. Libraries were sequenced on Illumina NovaSeq 6000 with paired-end 150 bp reads to a depth of approximately 5000 reads per cell.

Raw sequencing data were processed and read in alignment with the reference human genome (GRCh38/hg38), and gene expression matrices were generated by the 10X Genomics CellRanger v6,1. Quality control was performed using Seurat v4.0 in R v4.1. High mitochondrial content cells were removed from all clusters by removing cells with >20% of Unique Molecular Identifiers (UMI) mapped to mitochondrial genes and <200 unique genes. Data were normalized with SCTransform,^9^ principal component analysis (PCA) was used for dimensionality reduction (RunPCA), and cells were clustered using the Leiden algorithm with 15 principal components (FindNeighbors and FindClusters) in Seurat.^10^ Clusters were visualized using the Uniform Manifold Approximation and Projection (UMAP) algorithm.^11^ Differential expression analysis between clusters was conducted using the Wilcoxon rank-sum test with a false discovery rate (FDR) < 0.05 and log2 fold change > 1. Cell types were annotated based on the expression of known marker genes.

We employed two complementary approaches to quantify tumor cell heterogeneity. All tumor cells from humanized mice (n=3) and non-humanized mice (n=2) were pooled into their respective groups. For each group, we calculated the geometric center (centroid) in UMAP space and measured the Euclidean distance of each cell to its group centroid: d = √[(x - x̄)² + (y - ȳ)²] where (x, y) represents the UMAP coordinates of each cell and (x̄, ȳ) represents the mean coordinates of all cells in the group. This distance metric provides a quantitative measure of cellular dispersion, with larger distances indicating greater heterogeneity in transcriptional states.

To evaluate apoptosis in single-cell RNA sequencing data, we calculated an apoptosis score for each cell using a curated set of key apoptosis markers: BAX, BAK1, BCL2, BCL2L1, CASP9, CYCS (intrinsic pathway), FAS, FADD, CASP8, CASP10 (extrinsic pathway), CASP3, CASP7 (execution pathway), and additional markers including PARP1, APAF1, and BID. Expression values were obtained from the normalized SCT (Single-Cell Transform) assay data. For each apoptosis marker gene present in the dataset, we calculated Z-scores by scaling the expression values. The final apoptosis score for each cell was determined by taking the mean of these Z-scores across all marker genes, providing a standardized measure of apoptotic pathway activity at the single-cell level.

**Public single-cell data analysis**

Immune cell populations in GBM were extracted from the Extended GBmap dataset, which was public Single-cell RNA sequencing data that comprised 53 primary GBM and 29 recurrent GBM cases.^12^ The extraction was performed based on Annotation_level3. The following cellular populations were analyzed: AC-like astrocytes, B cells, CD4/CD8 T cells, dendritic cells (DCs), endothelial cells, mesenchymal-like cells, mast cells, monocytes, mural cells, natural killer (NK) cells, NPC-like cells, neurons, neutrophils, oligodendrocyte precursor cells (OPCs), OPC-like cells, oligodendrocytes, plasma cells, radial glial cells, tumor-associated macrophage(TAM)-blood-derived monocytes/macrophages (BDM), and TAM-microglia (MG). Cells identified as human CD45-positive comprised a mixture of B cells, CD4/CD8 T cells, DCs, mast cells, monocytes, NK cells, TAM-BDM, and TAM-MG. Subsequently, the proportion of each cell type was calculated relative to the total CD45-positive cell population.

The processed Seurat object file (CoreGBmap.rds) was obtained from the cellxgene website (Harmonized single-cell landscape, intercellular crosstalk and tumor architecture of glioblastoma, https://cellxgene.cziscience.com/collections/999f2a15-3d7e-440b-96ae-2c806799c08c). Subsequent analyses were performed using Seurat v5.1.0 and R v4.3.3. T cells were extracted based on the following information “original_celltype” annotations: “CD4+”, “CD8+”, “T cells”, “T cells 1”, “T cells 2”, “T cells 3”, “Treg”, and “Regulatory T cells” from 11 recurrent GBM patients. For a focused analysis of Tregs, cells annotated as “Treg” and “Regulatory T cells” were specifically extracted from the same 11 recurrent GBM patients. Macrophages and Monocytes were extracted based on the following information “annotation_level_4” annotations: “Mono anti-infl”, “Mono hypoxia”, “Mono naive”, “TAM-BDM IN”, “TAM-BDM MHC”, “TAM-BDM anti-infl” and “TAM-BDM hypoxia/MES” from 11 recurrent GBM patients. Data normalization was conducted using the ScaleData function in Seurat. Dimensionality reduction was achieved through PCA using the RunPCA function. Cell clustering was performed using the Leiden algorithm with appropriate principal components, implemented through the FindNeighbors and FindClusters functions in Seurat.^10^ The resulting clusters were visualized using the UMAP algorithm.^11^

**Statistical analysis**

Results were analyzed using a two-tailed Student’s t-test, 2-way analysis of variance (ANOVA), or the Mann-Whitney U‐test in GraphPad Prism 5.0 software or IBM SPSS Statistics 18 software to assess statistical significance. Kaplan-Meier survival analyses were performed using the log-rank test. p values <0.05 were statistically significant.

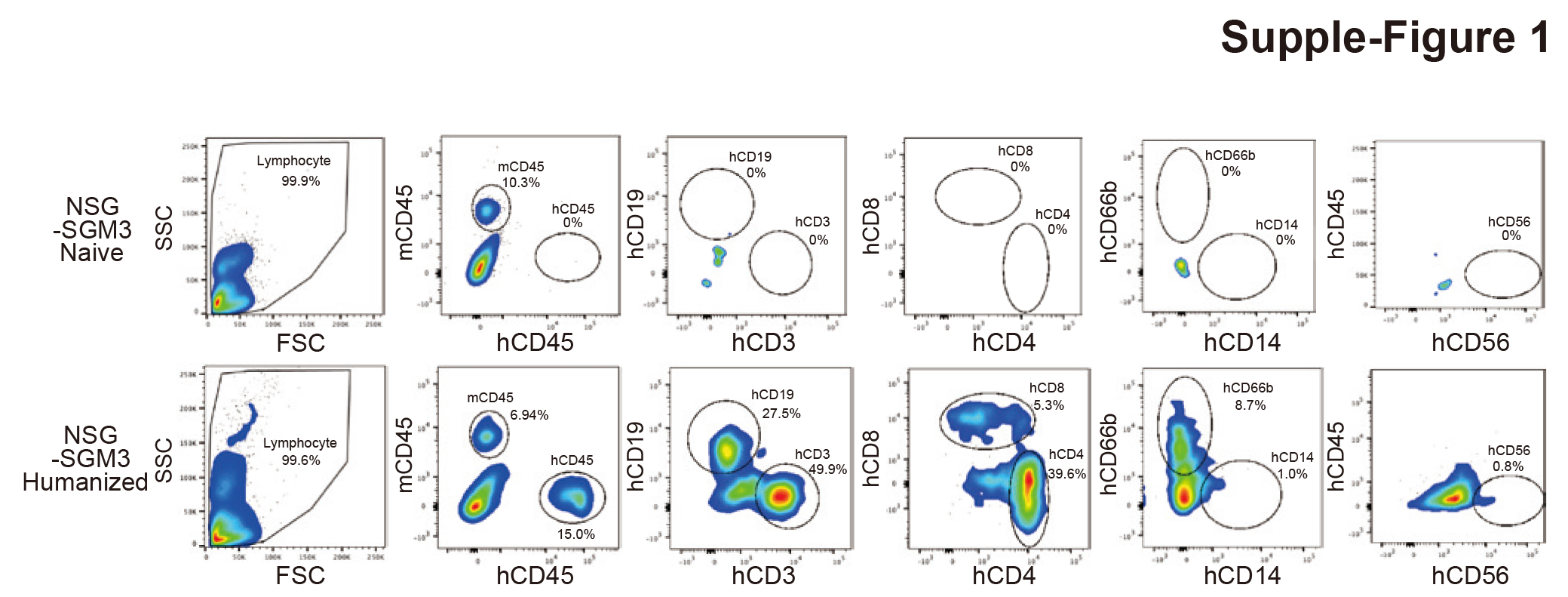
:

**Supplemental Figure 1**: **Reconstitution of human immune cells in humanized mice peripheral blood.** Representative data of Flow cytometry analysis of immune cells in the peripheral blood of humanized mice. %mouse CD45 (mCD45) and human CD45 (hCD45) in total cells. %hCD19, hCD3, hCD8, hCD4, hCD66b, hCD14 and hCD56 in hCD45 populations.

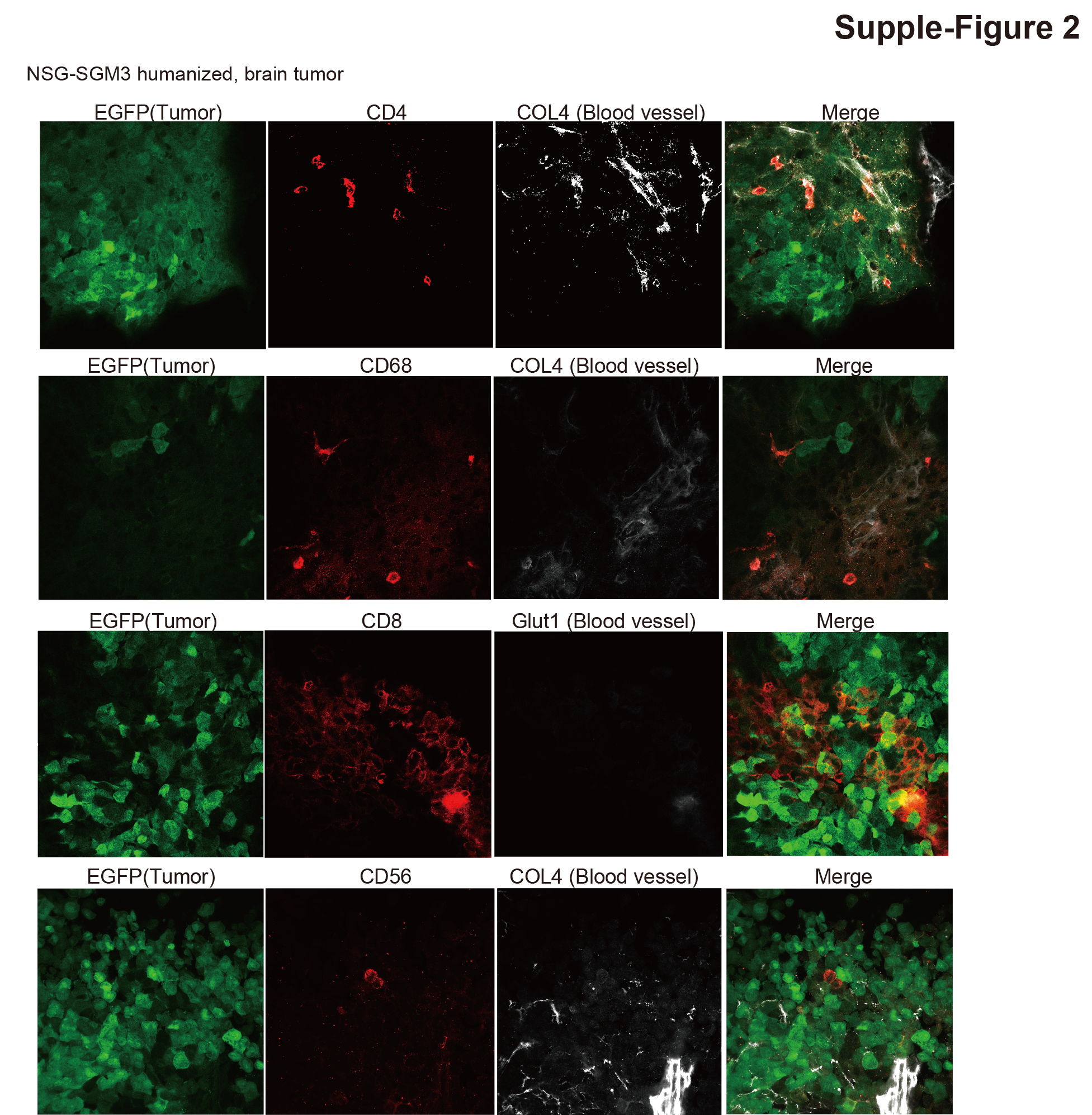

**Supplemental Figure 2**: **Human immune cells' localization into humanized mice's brain tumors.** Representative immunofluorescence staining of tissue sections from humanized mice glioma models. Tumor cells express EGFP (green). Blood vessels (white) were immunostained with antibodies against Collagen type IV (COL4) or Glut1. The expression sites of CD4, CD68, CD8, and CD56 are shown in red. Scale bars, 100 mm.

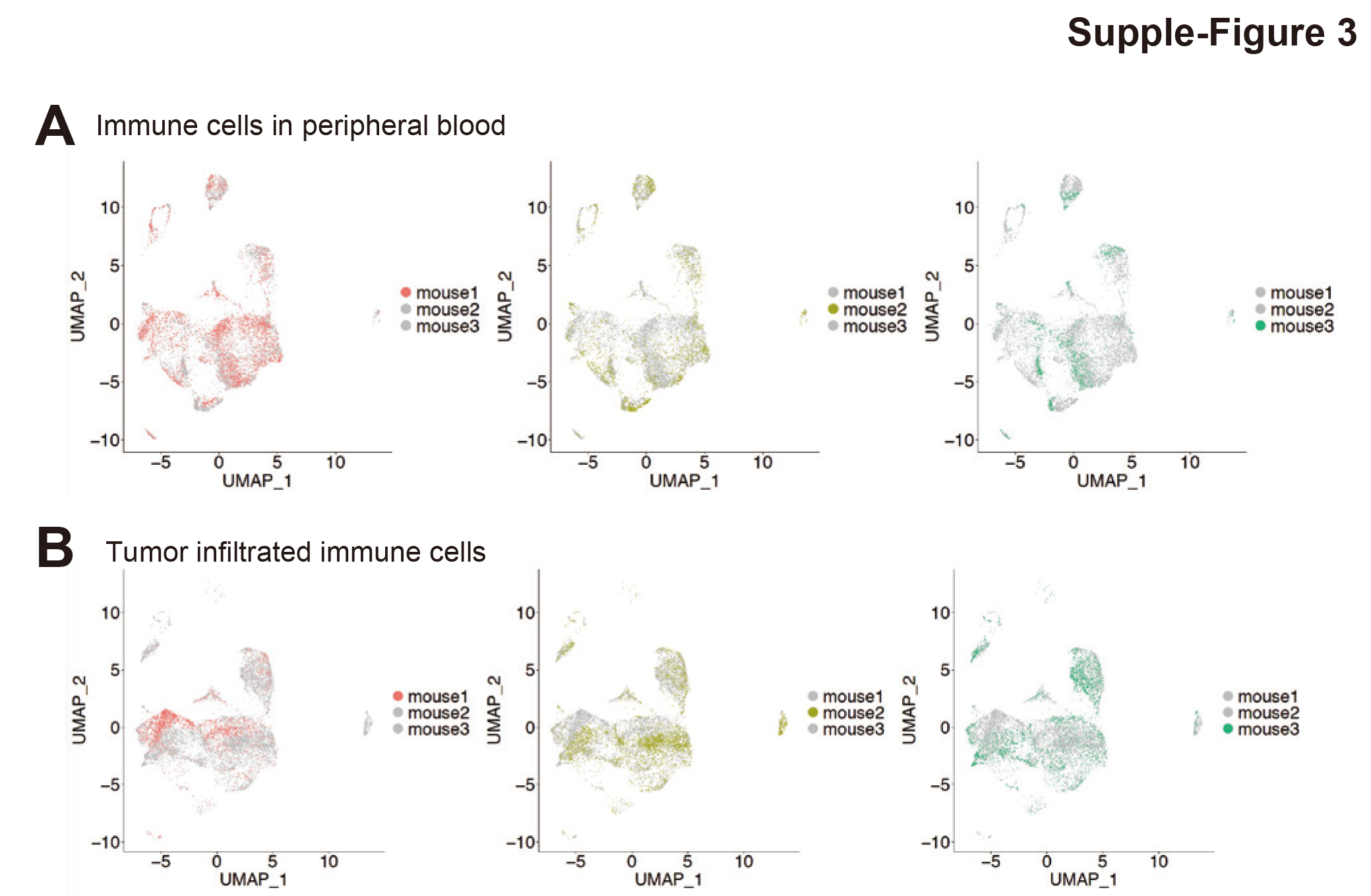
**Supplemental Figure 3**: **Single-cell RNA-seq analysis of CD45+ cells isolated from NOG-EXL humanized mice bearing transplanted JX14P-RT human glioblastoma cells. (**A) UMAP plot of CD45+ cells in peripheral blood (PBs) and (B) tumor-infiltrating lymphocytes (TILs) from three humanized mice.

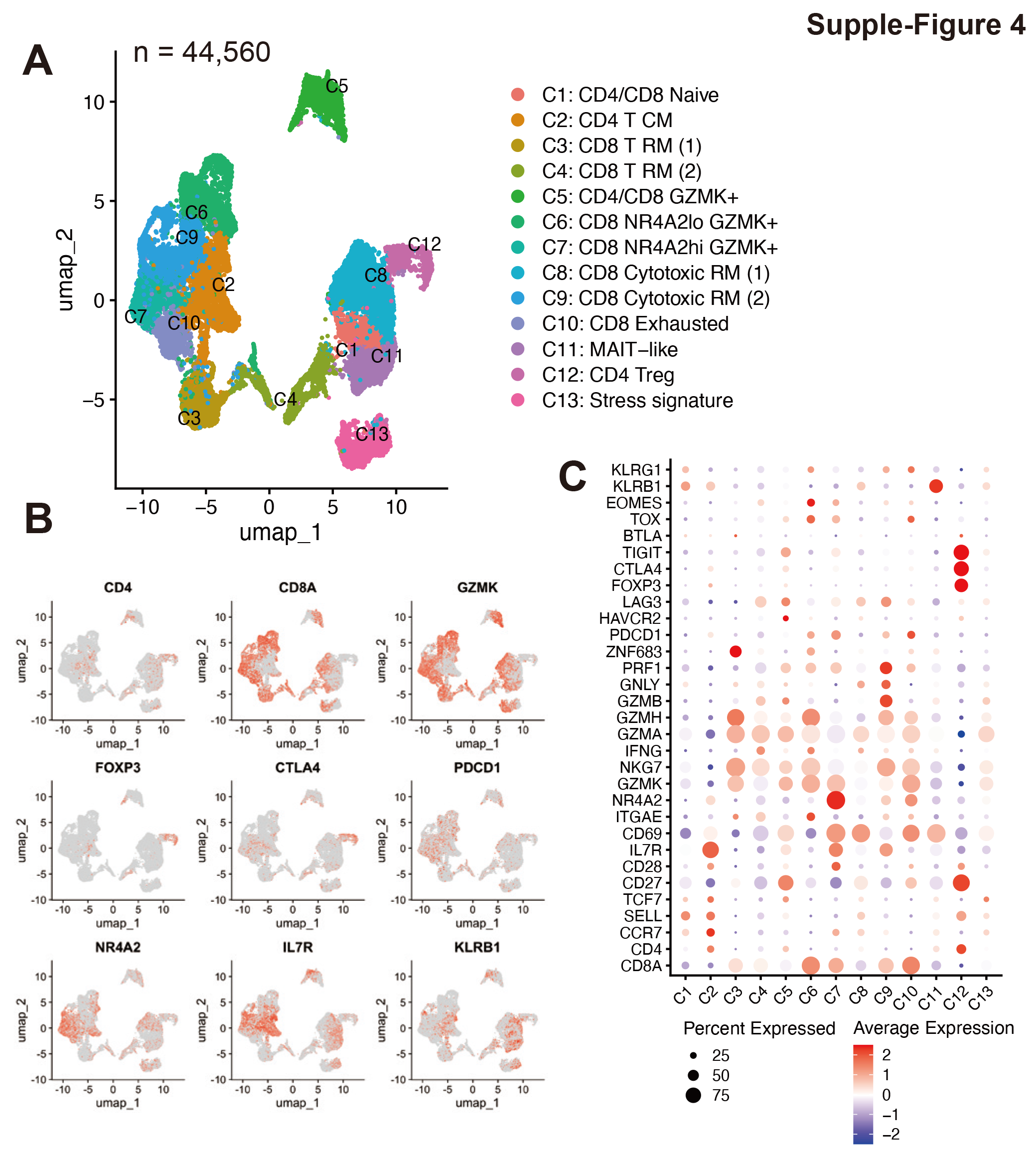

**Supplemental Figure 4**: **Single-cell RNA-seq analysis of T cell subsets in human recurrent GBM from public dataset.** (A) UMAP plot of T cell populations (n = 44,560 cells) showing 13 distinct cell types identified in the tumor microenvironment. (B) Feature plot showing the expression of key T cell markers across the identified subsets. The intensity of color represents the level of gene expression for KLRB1, PDCD1 (PD-1), IL7R, CTLA4, CD8A, NR4A2, FOXP3, CD4, and GZMK. (C) Heatmap displaying the expression patterns of T cell-associated genes across the 13 identified subsets.

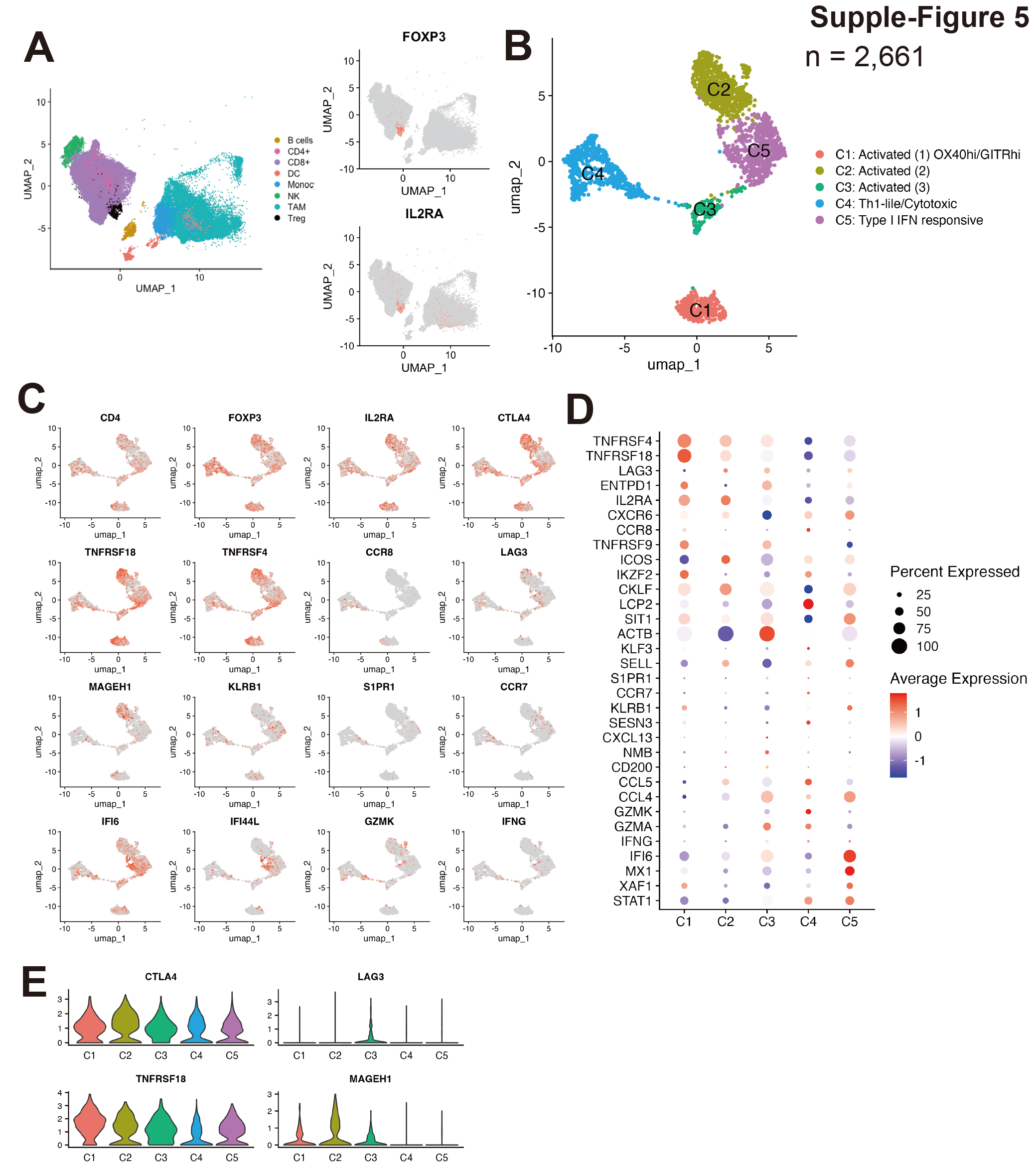
**Supplemental Figure 5**: **Single-cell RNA-seq analysis of regulatory T cell subsets in human recurrent GBM. (**A) UMAP plot of TILs showing Treg populations (black dots) with high FOXP3 and IL2RA (CD25) gene expression. (B) UMAP plot of Treg cells (n = 2,661) showing five cell types. (C) Feature plots displaying the expression of individual genes across the Treg population. (D) Heatmap showing differential gene expression across the identified Treg subsets. (E) Violin plot showing the expression of four key immune checkpoint molecules (CTLA4, TIGIT, PD1, and LAG3) across the T cell subsets.

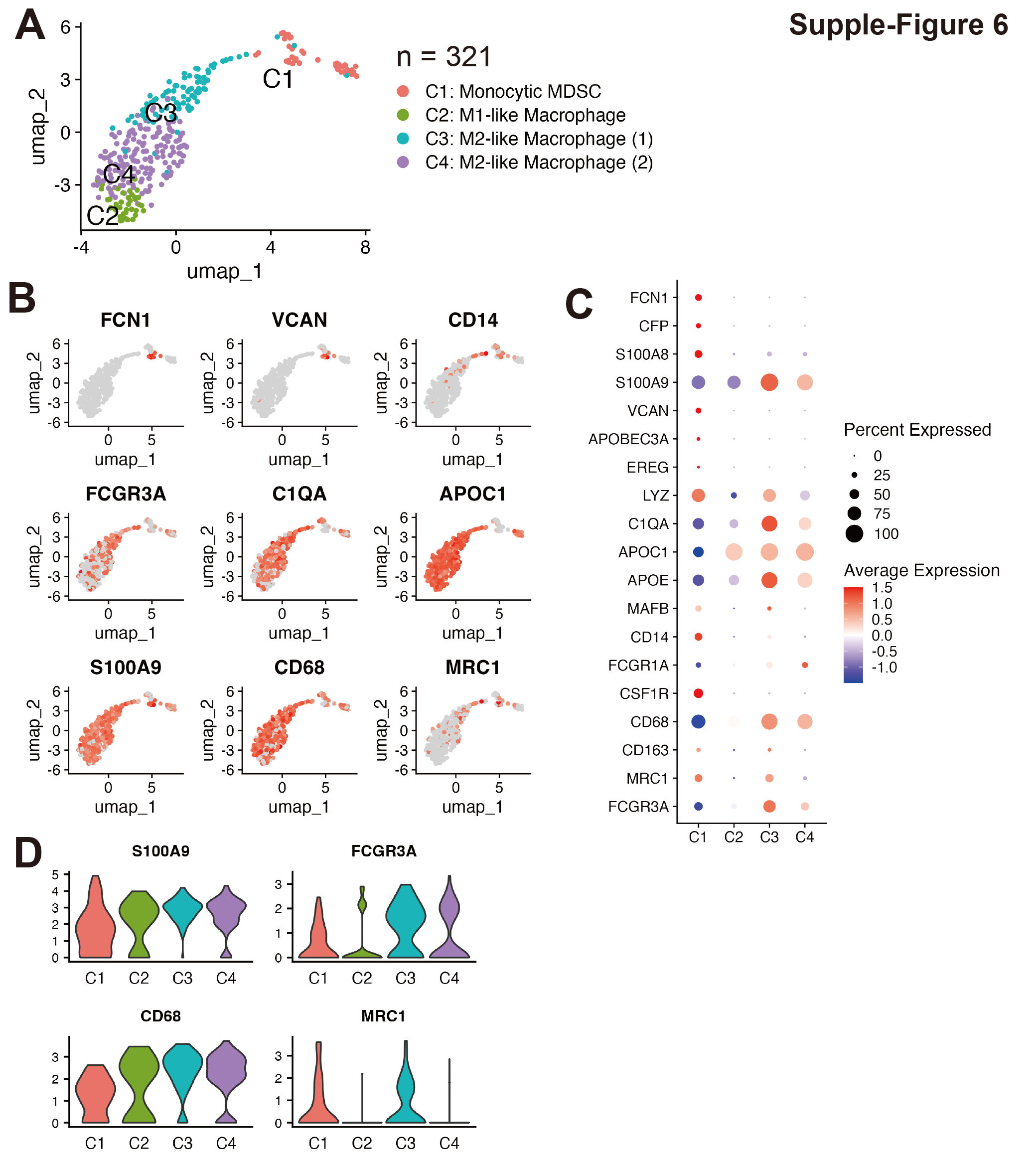

**Supplemental Figure 6**: **Subcluster analysis of macrophage and monocyte subpopulation**. (A) UMAP plot of M-MDSC, macrophage, and monocyte populations (n = 321 cells) showing four distinct cell types identified in the tumor microenvironment. (B) Feature plot showing the expression of key markers across the identified subsets. The intensity of color represents the level of gene expression for FCN1, VCAN, CD14, FCGR3A, C1QA, APOC1, S100A9, CD68 and MRC1 (CD206). (C) Heatmap displaying the expression patterns of macrophages and monocyte genes across the four identified subsets. (D) Dot plot showing the subsets’ expression of four key molecules (S100A9, FCGR3A, CD68, and MRC1).**:**

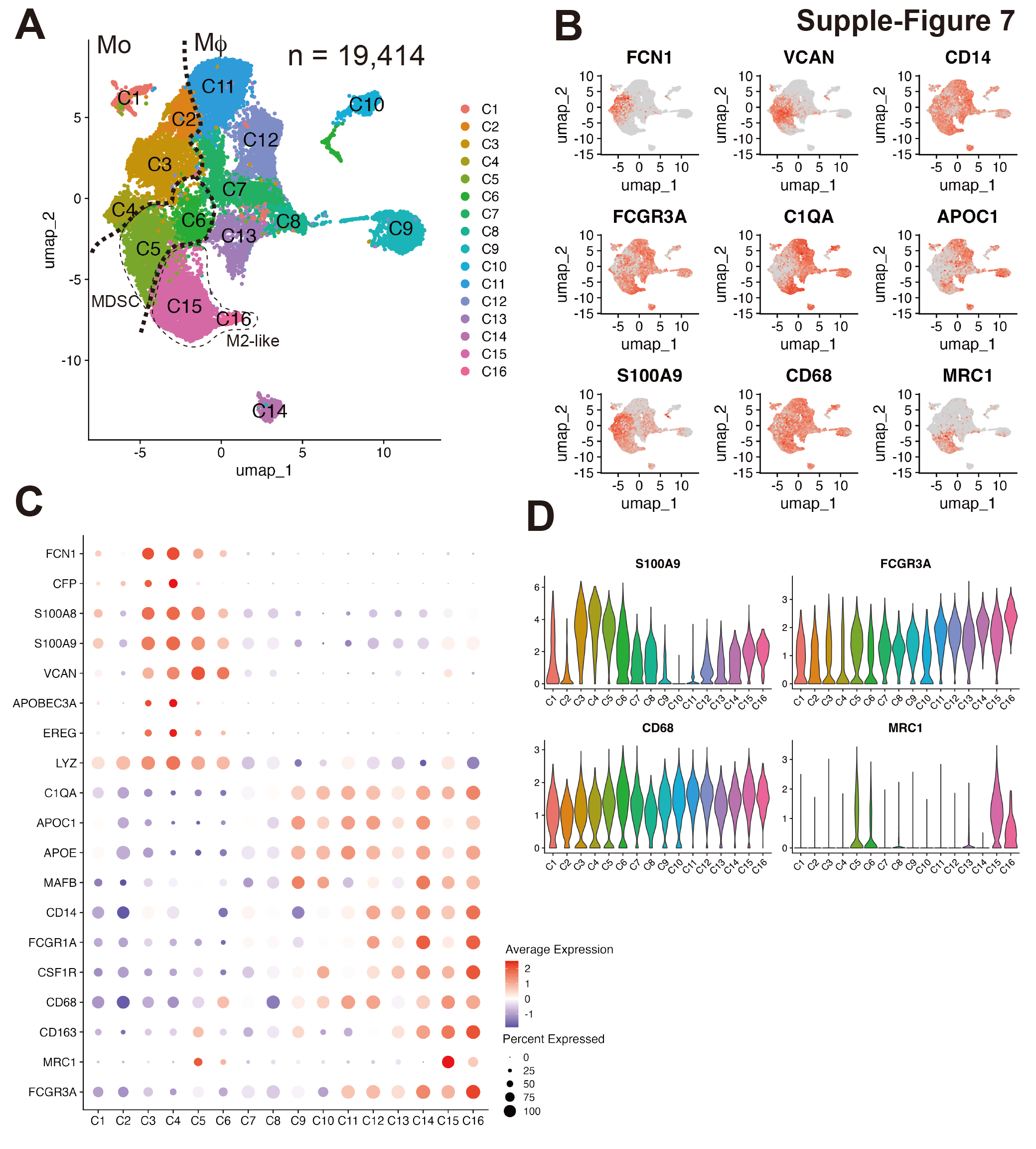
**Supplemental Figure 7**: **Single-cell RNA-seq analysis of macrophage and monocyte in human recurrent GBM.** (A) UMAP plot of M-MDSC, macrophage, and monocyte populations (n = 19,414 cells) showing 16 distinct cell types identified in the tumor microenvironment. (B) Feature plot showing the expression of key markers across the identified subsets. The intensity of color represents the level of gene expression for FCN1, VCAN, CD14, FCGR3A, C1QA, APOC1, S100A9, CD68 and MRC1 (CD206). (C) Heatmap displaying the expression patterns of macrophages and monocyte genes across the 16 identified subsets. (D) Dot plot showing the subsets' expression of four key molecules (S100A9, FCGR3A, CD68, and MRC1).

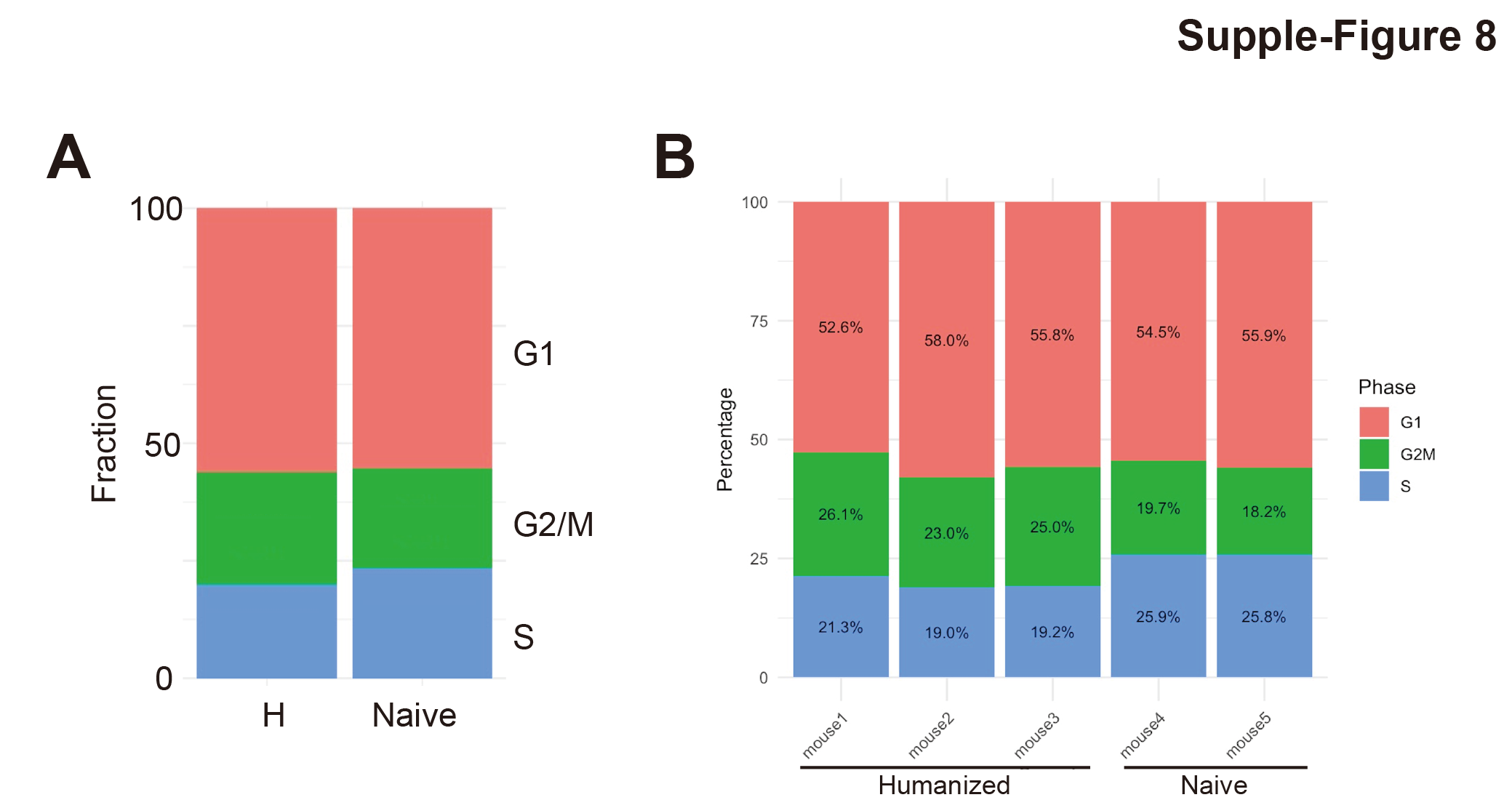

**Supplemental Figure 8**: **Cell cycle genes between humanized and naive mouse models**. (A) Fraction of tumor cells in different cell cycle phases (G1, G2/M, S) for humanized and naive mice. (B) Bar plots showing the state of the cell cycle of tumor cells from individual humanized mice (n=3) and naive mice (n=2).

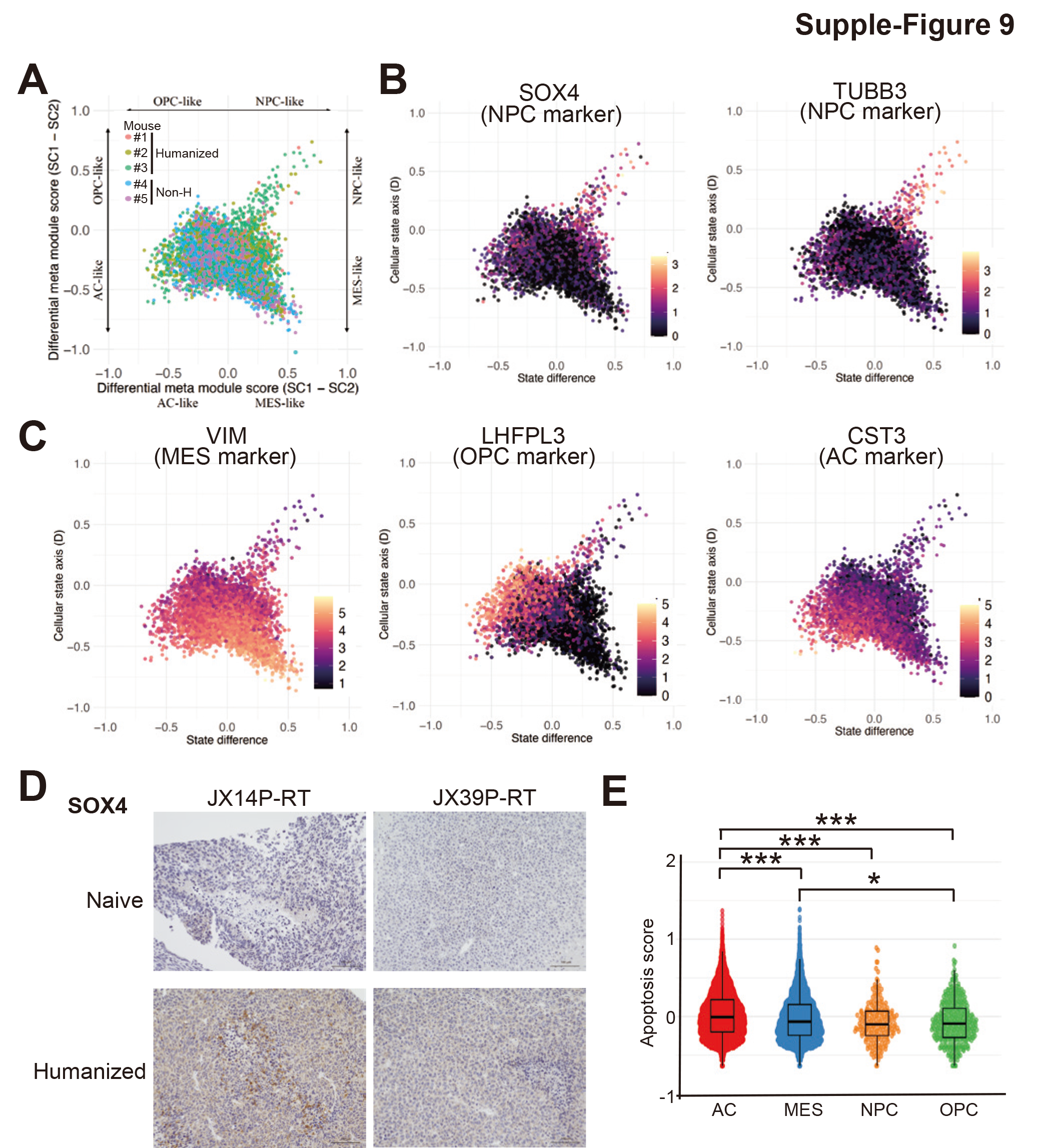

**Supplemental Figure 9**: **Cellular states between humanized and naive mouse models.** (A) UMAP plots of the cell state of tumor cells from individual humanized mice (n=3) and naive mice (n=2). (B) Feature plot showing the expression of key cell state markers of neural progenitor (NPC). The intensity of color represents the level of gene expression for SOX4 and TUBB3. (C) Plot showing the expression of key cell state markers of mesenchymal (MES), oligodendrocyte-progenitor (OPC), and astrocyte (AC). The intensity of color shows the level of gene expression for VIM, LHFPL3, and CST3. (D) Representative immunohistochemistry staining of SOX4 in humanized mice and naïve mice. (E) Distribution of apoptosis scores among tumor cells within AC, MES, NPC, and OPC cellular states. Kruskal-Wallis rank sum test, *P < 0.05, ***P < 0.001.

Table 1.

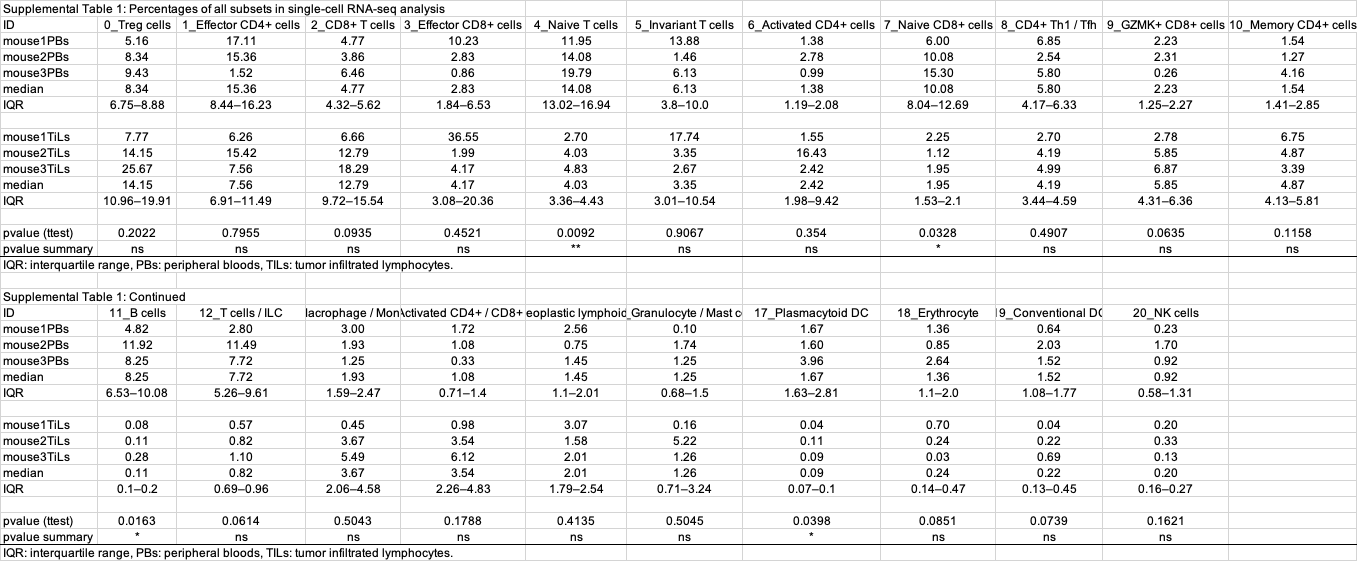

Table. 2
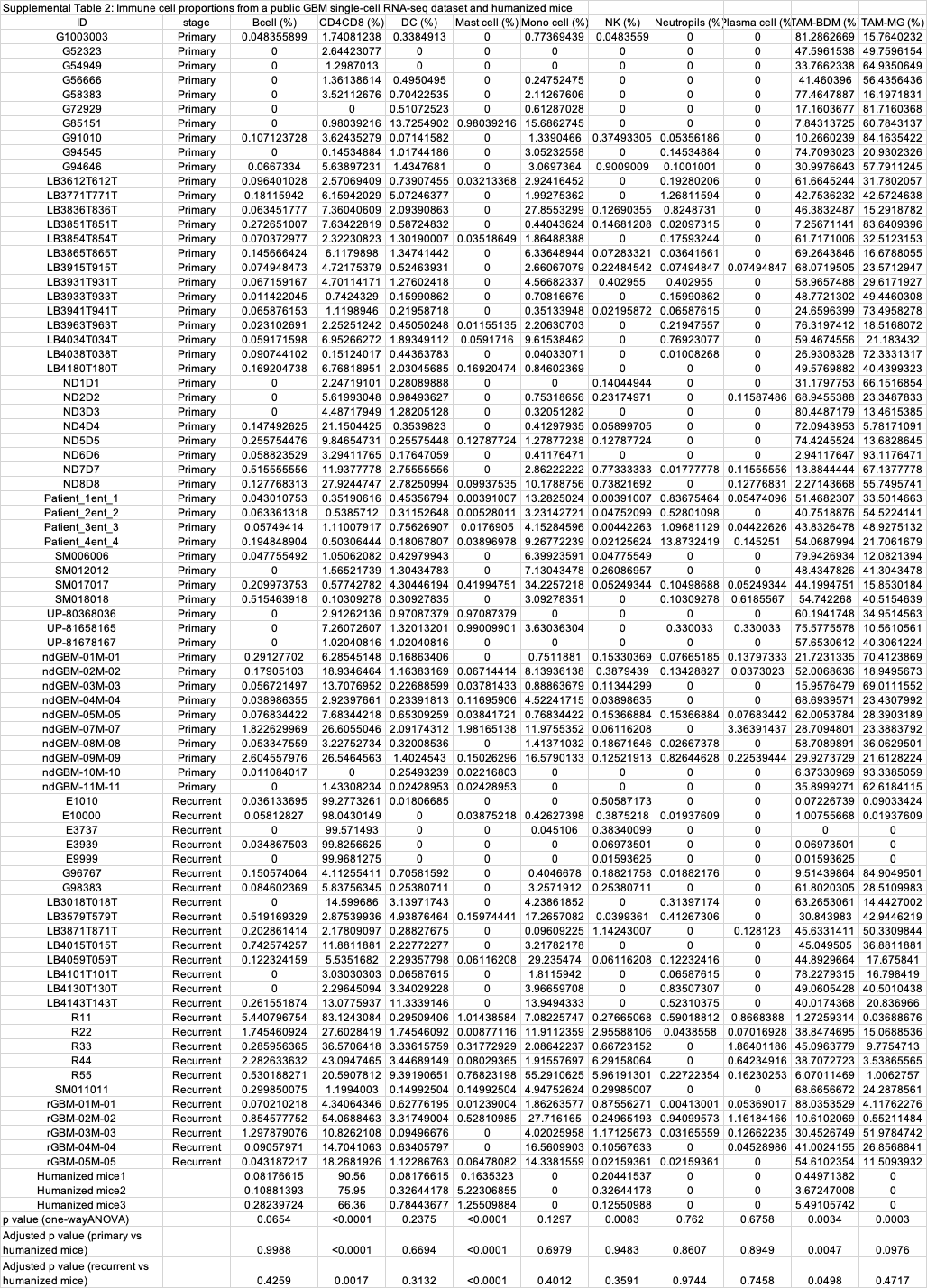
